# Signalling Network of Breast Cancer Cells in Response to Progesterone

**DOI:** 10.1101/2020.11.03.366401

**Authors:** Roni H. G. Wright, Viviana Vastolo, Javier Quilez Oliete, José Carbonell-Caballero, Miguel Beato

## Abstract

Breast cancer cells enter into the cell cycle following progestin exposure by the activation of signalling cascades involving a plethora of enzymes, transcription factors and co-factors that transmit the external signal from the cell membrane to chromatin, ultimately leading to a change of the gene expression program. Although many of the events within the signalling network have been described in isolation, how they globally team up to generate the final cell response is unclear. In this study we use antibody microarrays and phosphoproteomics to provide a dynamic global signalling map that reveals new key regulated proteins and links between previously known and novel pathways. Detailed analysis of the data revealed intriguing changes in protein complexes involved in nuclear structure, EMT, cell adhesion, as well as transcription factors previously not associated with breast cancer proliferation. As different post-translational modifications can mediate complex crosstalk mechanisms and massive PARylation is also rapidly induced by progestins, we provide details of important chromatin regulatory complexes containing both phosphorylated and PARylated proteins. This study contributes an important resource for the scientific community, as it identifies novel players and connections meaningful for breast cancer cell biology and potentially relevant for cancer management. (197 w)

## Background

Female steroid hormones, oestrogen and progesterone play a key role not only in the normal development of target tissues during puberty, pregnancy and menopause but also in breast and endometrium cancer cell proliferation. Breast cancer cells respond to progestin exposure with two intermingled pathways that culminate in extensive gene expression changes and entry in the cell cycle. The classical view is that the hormone diffuses through the cell membrane and binds to intracellular progesterone receptors (PR), which are maintained in an inactive state by a chaperone complex, including Heat Shock Proteins 70 and 90 (HSP70/90). Upon hormone binding, PR weakens its interaction with the chaperones, dimerizes and moves to chromatin where eventually binds to palindromic DNA sequences called progesterone responsive elements (PREs) (Pina et al., 1990). Once bound to chromatin, PR recruits various co-regulators and chromatin remodellers that modulate access for the transcription machinery including RNA polymerase II (Beato et al., 1995).

This simplified model was completed later by the finding that a tiny fraction of PR (3-5%) is attached to the cell membrane via palmitoylation at C820 (Migliaccio et al., 1998; Pedram et al., 2007), forming a complex with estrogen receptor alpha (ERa) (Ballare et al., 2003). Upon binding progestins, the membrane anchored PR activates SRC, either directly (Boonyaratanakornkit et al., 2001) or via ERa, initiating a kinase signalling pathway that ends in activation of the extracellular signal-regulated kinase (ERK) (Ballare et al., 2003). ERK1 phosphorylates intracellular PR at S294 favouring its dissociation from the chaperone complex. In the cell nucleus ERK1 activates MSK1 (Mitogen-and stress activated protein kinase 1), resulting in the formation of a ternary complex of PR-ERK1-MSK1, which is the active form of PR able to regulate chromatin structure and gene expression. ERK also activates cyclin dependent kinase 2 (CDK2) that in turns activates ARTD1 (ADP-ribose transferase 1) by phosphorylating two serines in the NAD+ binding pocket (Wright et al., 2012). Phosphorylation contributes to dissociation of histone H1 and H2A/H2B dimers (Vicent et al., 2006; Wright et al., 2012) and to local chromatin opening by further recruitment of transcription factors, co-regulators, histone modifiers (PCA, P3000) and ATP-dependent chromatin remodellers (NURF and BAF), ultimately leading to the activation of gene expression changes (Vicent et al., 2011; Vicent et al., 2009a; Vicent et al., 2010). Moreover, there is also evidence for the activation by progesterone of other signaling pathways induced by progesterone, such as AKT (Fu et al., 2010), cAMP (Garg et al., 2017; Takahashi et al., 2009), GSK3 (Rider et al., 2006) and STAT (Hagan et al., 2013). However, many of these studies use different cells of a different types of cells derived from endometrial or ovarian tissues (Lee and Kim, 2014; Wang et al., 2007).

In addition to the key role of the kinase cascades, we have identified a pivotal role for another post-translational modification in progestin induced gene regulation; namely Poly-ADP-ribosylation (PARylation). As discussed above the PAR polymerase PARP1, also known as ADP-ribosyltransferase 1 (ARTD1), is activated within the initial minutes following hormone exposure via phosphorylation by CDK2 (Wright et al., 2012), giving rise to a large increase in PARylation within the cell nucleus (Wright et al., 2012). Parylation of ARTD1 itself and of chromatin proteins is essential for the initial dissociation of histone H1 (Nacht et al., 2016; Vicent et al., 2016). We also found that degradation of PAR to ADP-Ribose by PAR glycohydrolase (PARG) is required for complete chromatin remodelling and activation of the gene expression network (Wright et al., 2016). Mass spec analysis of the proteins interacting with PAR in T47D cells exposed to progestins revealed structural proteins, DNA damage response proteins and chromatin modifying enzymes (Wright et al., 2016). One key enzyme identified in this study was NUDT5 or NUDIX5 (Nudix hydrolase 5), which hydrolyses ADPR to AMP and ribose-5-phosphate. Subsequently, we found that upon dephosphorylation at T45, NUDT5 can use ADPR and diphosphate for the synthesis of ATP (Wright et al., 2016). In this way, part of the ATP consumed during the synthesis of NAD^+^ and stored in PAR is recovered and used for chromatin remodeling and changes in gene expression. The synthesis of nuclear ATP is transient, peaking at 40 minutes after hormone exposure and returning to basal levels after ~60 min. However, although we know that nuclear ATP synthesis is essential for the initial chromatin remodeling, the role of the nuclear ATP at later time points is unclear. We can envision several hypotheses; as a direct, local source of ATP for the massive amount of ATP-dependent chromatin remodelling and 3D conformational changes induced by hormone (Le Dily et al., 2019; Vicent et al., 2011) or to facilitate phase separation of chromatin fiber (Wright et al., 2019). In any case, we know that nuclear ATP synthesis by NUDT5 is essential for the generation and maintenance of the cancer stem cell population (Pickup et al., 2019).

Over the past years, there has been a large number of studies investigating the role and mechanism of action of one or more of the pathway components in response to progesterone exposure, revealing a dynamic crosstalk between canonical pathways (Boonyaratanakornkit et al., 2008; Faivre et al., 2005; Qiu et al., 2003; Skildum et al., 2005). However, these studies focused on one or very few components at a time, and do not explain how these various pathways interact and coordinate the cell response to hormone. The work described here aims to provide a more comprehensive map of progesterone signalling in breast cancer cells, combining antibody arrays technology, shotgun proteomics, and previously published PARylation datasets to develop for the first-time a global map of the dynamic signalling events induced by progestins in breast cancer cells.

## Results

### 1. Prior Knowledge Network

Before starting to add quantitative dynamic data to the already existing knowledge of progesterone signalling events in breast cancer cells, we have generated a “prior knowledge network (PKN)”, based on the published literature (Fig.S1A-B, and Supplementary material; Cytoscape Session 1). Each protein-protein interaction is characterized based on type (interaction, phosphorylation or dissociation) and is displayed as a unique edge. The corresponding literature is given in Fig. S1B and within the Cytoscape session. The PKN already shows the key role played by kinases in the response of breast cancer cells to progestins. First, progestins via ERa activate SRC1 that phosphorylates MAPKK1, that activates ERK1, that phosphorylates PGR, resulting in dissociation from the HSP90A and B proteins (Haverinen et al., 2001; Smith, 1993). Activated ERK1 also phosphorylates ERa at S118 (Kato et al., 1995). ERK1 in association with hormone receptors translocates to the cell nucleus where it phosphorylates MSK1 (Reyes et al 2016), leading to the formation of an active complex PR-ERK-MSK1 that interacts with chromatin containing accessible PRE. Activated PR also interacts with PLK1 that activates MLL2 (Wierer et al., 2013), with CDK2 that phosphorylates and activates ARTD1 (Wright et al., 2012), and with JAK2 that activates STAT5 (Hagan et al., 2013). Simultaneously, membrane activated SRC1, also activate RAS and EGFR (Boonyaratanakornkit et al., 2007), which feeds back activating the MAPK cascade. Membrane associated ERa also activates PI3K and cAMP, which upon binding with AKT and PKA respectively lead to the activation (via interaction and direct phosphorylation) of GSK3, mTOR (Ciruelos Gil, 2014; Ortega et al., 2020) and the arginine methyltransferases CARM1 and PRMT1 within the nucleus (Lange, 2008; Li et al., 2003; Malbeteau et al., 2020). This brief description of the PKN shows that it already encompasses a great degree of complexity and complementary connection that need additional data to be resolved.

### 2. Microarrays of antibodies to phosphorylated sites in proteins

Our plan was to combine antibody microarray technology and shotgun phosphoproteomics in T47D^M^ breast cancer cells exposed to 10 nM R5020 for different lengths of time, as previously described (Wright et al., 2012). For each experimental approach and exposure time total protein extracts were harvested in triplicate. For the antibody arrays, data was collected, filtered for quality control and summarized as log_2_ ratio over time zero (as described in materials and methods, Fig. S2A). This dataset provided 246 unique phosphorylation sites corresponding to 155 proteins (Fig. 1A, Supplementary Table S1). The majority of proteins contain 1 phosphorylation site, although for several proteins (Tau, RB1, MAP2K1, PTK2 and the protein kinase RPS6KA1) 7 or more significantly regulated phosphorylation sites were identified (Fig. S2B).

**Fig. 1.**
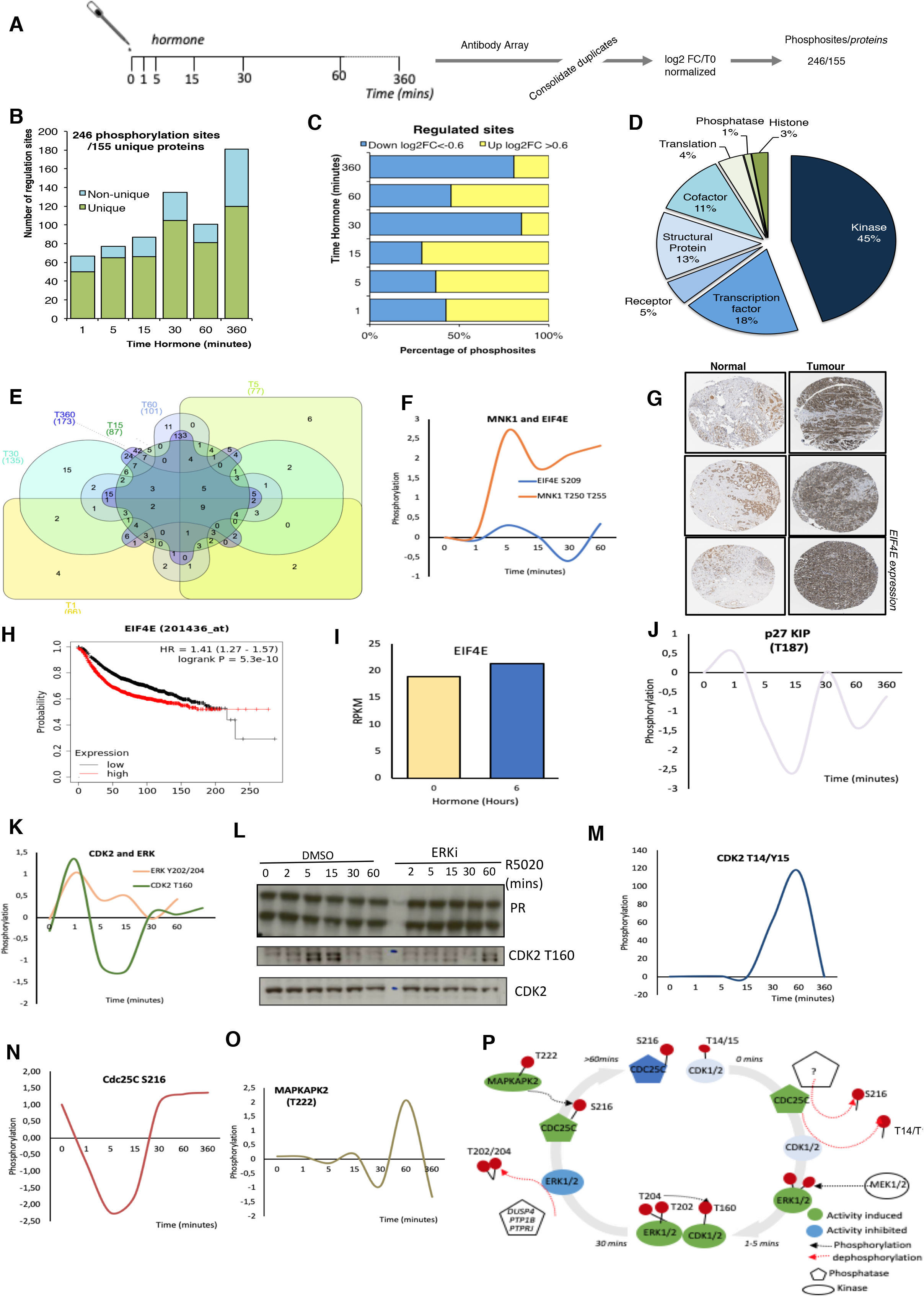
Targeted Antibody Array Phosphorylation data following hormone. **A)** Schematic overview of experimental procedure. Synchronised T47D cells were exposed to hormone for the length of time indicated. Triplicate samples were harvested and phosphorylated proteins were identified using an antibody microarray (see materials and methods). Data was log_2_ normalised resulting in a total of 246 significant phosphosites from 155 unique proteins. **B)** Number of regulated sites per time point, log_2_FC (Fold change) >0.6<-0.6 versus time 0. **C)** Breakdown of up (>0.6log_2_FC) and down (<-0.6 log_2_ FC) per time point versus T0. **D)** Functional classification of the proteins identified as significantly phosphorylated across all time points, individual time point functional analysis (per time point see Fig. S2C). **E)** Venn diagram showing the overlap of significantly regulated phosphorylation sites across all time points. **F)** Log_2_ FC following hormone of phosphorylated Mnk1 (T197/202) and EIF4E S209. **G)** Expression of EIF4E in normal versus breast tumour samples from breast cancer patients (Protein Atlas, see materials and methods). **H)** Kaplan Meyer overall survival stratifying patients based on the expression level of EIF4E in breast cancer data set (p=5.3e-10). **I)** mRNA expression level of EIF4E in T47D cells treated with hormone. Dynamics of **(J)** p27/KIP T187 **(K)** CDK2 T160, ERK Y202/204. **L).** Expression level of total PR and CDK2 and phospho-CDK2 T160 in T47D breast cancer cells exposed to hormone in the presence or absence of ERK inhibitor (ERKii) as determined by western blotting using specific antibodies. **(M)** CDK2 T14/Y15 phosphorylation in response to hormone as determined by antibody array. Dynamics of **(N)** Cdc25C S216 and **(O)** MAPKAPK2 T222 phosphorylation in response to hormone as determined by antibody array. **P)**. Model for CDK2/ERK dynamic activation and deactivation in response to hormone based on the data presented in Fig. 1J-O.

Analysis of the number of phosphorylation sites clearly showed a rapid activation already 1-minute following hormone (68 significant phosphorylation events Fig 1B). Signalling persists throughout the time course showing two peaks at 30- and 360-minutes following hormone (Fig. 1B). Phosphorylation sites were characterized as up or down-regulated, using a threshold for the log_2_ fold change with respect to time 0 of −0.6< or >0.6 respectively (Fig 1C). We see a trend for early phosphorylation sites to be dynamically increased compared to time zero, in contrast to later time points where protein phosphorylation sites as a whole decrease compared to time zero (Fig 1C). The majority of phosphorylation sites identified belong to protein kinases (45%), co-factors (11%), transcription factors (18%) and structural proteins (13%). (Fig 1D). Combining the identified phosphorylation sites over the time course reveals that the majority of sites are regulated at more than one time point (Fig 1E), however the protein function enrichment does not alter significantly over time, with kinases and transcription factors being the main protein groups where the phosphorylation sites are observed (Fig. S2C).

Pathway and gene ontology (GO) for biological function (BP) molecular function (MF) analysis revealed a significant increase in Cancer pathways (Fig S2D), signal transduction, biopolymer metabolic process and kinase activity (Fig 2SE and F). Within this dataset we observed a strongly upregulated phosphorylation of the MAPK Signal-Integrating Kinase 1, MNK1 at T250/T255 (Fig. 1F) in T47D in response to progesterone stimulation. Phosphorylation of MNK1 at T250/T255 by ERK induces the activity of MNK1 (Dolniak et al., 2008). Once activated, MNK1 phosphorylates its targets, including the proto-oncogene Eukaryotic Translation Initiation Factor 4E (EIF4E), for which we also observed a modest phosphorylation which follows a similar pattern to MNK1 (Fig. 1F). Activation of MNK1 has been shown to promote cell proliferation thus MNK1 inhibitors appear as an exciting opportunity for cancer therapy. MNK1 signalling play a key role in invasive breast cancer growth (Guo et al., 2019), MNK1 inhibitors have been shown to block breast cancer proliferation in multiple cell lines (Wheater et al., 2010), and its downstream target EIF4E is 8 overexpressed in tumour versus normal samples from breast cancer patients (Fig. 1G) and associated with a poor overall survival (Fig. 1H). Our results are the first indication that MNK1 activation may be relevant for progesterone induced breast cancer cell proliferation.

We found that CDK2 plays an important role in progesterone signaling, activating ARTD1, and phosphorylating histone H1 (Wright et al., 2012). CDK2 activity is controlled by the formation of an active complex with the cyclin partner; either Cyclin E or A. In addition to binding the cyclin partner, CDKs are also controlled via interactions with Kinase Inhibitory Proteins (KIPs). p27/KIP is rapidly dephosphorylated at T187 in response to hormone, dropping sharply at 1 minute after hormone exposure (Fig. 1J), when CDK2 is phosphorylated and activated. Phosphorylation of p27 at T187 results in the proteins ubiquitination and degradation and inhibits the interaction with CDK2 (Grimmler et al., 2007), which would result in the release of CDK2 from the inhibitory protein resulting in the activation of ARTD1 and subsequent nuclear effects. In addition, we observe the coordinated activation of the upstream kinase of CDK2 at T160; ERK at Y202/204 (Fig. 1K) and could validate the phosphorylation of CDK2 T160 via ERK by western blotting in the presence of ERK inhibition (Fig. 1L). CDK2 at T160 is the active phosphorylation site of CDK2 peaking at 1-minute following hormone exposure (Fig. 1K) in contrast to the inactive phosphorylation site of CDK2 (T14/Y15) which peaks at 60 minutes following hormone exposure to silence the kinase (Fig. 1M). The activity of the phosphatase CDC25C is key for the removal of the inhibitory T14/Y15 phosphorylation sites of CDK2. The phosphatase itself is inactivated by phosphorylation at S216. We observe a peak in CDC25C phosphorylation prior to and following 60 minutes of hormone exposure, which would permit the phosphorylation of the inhibitory phosphorylation site in CDK2 (Fig. 1M and N). Going one step further; MAPKAPK2, the kinase which phosphorylates CDC25C at S216 is activated following the same time dynamic as its target (Fig. 1O). Although the importance of CDK2 in progestin induced cell proliferation has been studied (Trevino et al., 2016; Wright et al., 2012) the complex mechanism of CDK2 activation; phosphorylation of active/inactive marks, activation and regulation of upstream phosphatases and kinases was not clear until now (Fig. 1P). These examples of the dynamic phosphorylation of MNK1 and CDK2 highlight the insight that can be gained by this type of global signaling datasets.

### 3. Shotgun phosphoproteomics

To complement the microarray dataset, we performed shotgun phosphoproteomic analysis using mass spec. Phospho-peptides from T47D^M^ cells exposed to 10 nM R5020 for the same duration as in the array experiments, were enriched using TiO2 and phosphorylated peptides identified by LC-MS-MS (Fig. 2A, Fig. S3A, Supplementary Table S1). We identified changes in 310 unique phosphorylation sites within 264 unique proteins (Fig 2B and C). The majority of proteins exhibited regulation of a single phosphosite, except for the serine/arginine repetitive matrix protein, SRRM1, involved in mRNA processing and the TP53 enhancing protein TP53BP1, that exhibited 8 and 10 regulated phosphorylation sites respectively (Fig. S3B). Most phosphorylation sites identified were phosphor-serine consistent with the biological ratio of residue specific phosphorylation (Fig 2D). Over the time course, changes at each time point were identified as either up (log_2_FC>0.6) or down (log_2_FC<-0.6) regulated (Fig. 2E). Up-regulated sites prevailed at early time points and many of these phosphorylation sites were significantly regulated at more than one time point (Fig. 2F). Pathway and GO-BP (Biological Process) and MF (Molecular Function) enrichment analysis was consistent with the antibody array enrichment and revealed an increase in pathways in cancer, biopolymer metabolic process and kinase activity (Fig. S3C-E).

**Fig. 2.**
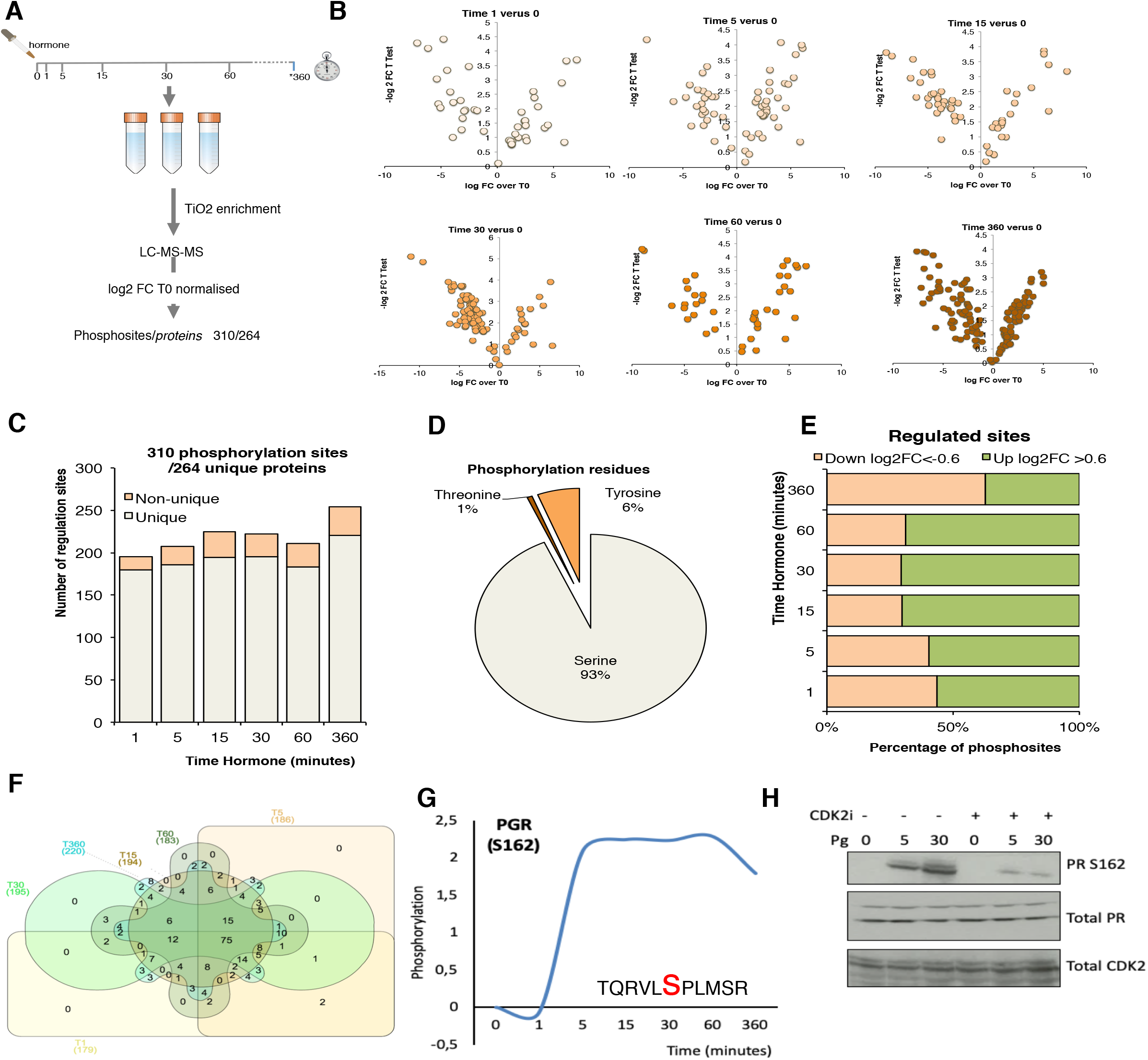
Phosphosite enriched Shotgun Proteomics following hormone. **A)** T47D cells were treated with hormone at the times indicated. Biological triplicates were enriched for phosphopeptides using TiO2 followed by LC-MS-MS peptide identification. Data was log_2_ normalised resulting in a total of 310 phosphosites from 264 unique proteins. **B)** Volcano plots showing phosphopeptide log_2_FC versus p-value for each of the time points following hormone. **C)** Number of significant phosphosites identified per time point. **D)** Analysis of the proportion of threonine, tyrosine and serine phosphorylated residues identified. **E)** Breakdown of up (>0.6log_2_FC) and down (<-0.6 log_2_ FC) per time point versus T0. **F)** Venn diagram showing the overlap of significantly regulated phosphorylation sites over time. **G)** Phosphorylation of progesterone receptor PR (S162), following progesterone validated by western blotting in presence or absence of CDK2 inhibitor **(H).** Total PR levels are shown as a loading control.

PR S294 is rapidly phosphorylated in response to hormone resulting in its activation and dissociation from chaperone complexes and increase protein turnover (Lange et al., 2000). In recent years it has been shown that clinical samples assigned as “PR low” actually have elevated levels of phosphorylated PR S294 and that this phosphorylation is associated with a genetic signature linked to cancer stem cell growth and increased recurrence which may have implications for the treatment of PR low patients with anti-progestins (Knutson et al., 2017). Phosphorylation of PR S162 in the hinge region showed a strong hormone induced increase by mass spec (Fig. 2G and H). Phosphorylation within this region of PGR has been previously reported to be mediated by CDK2 (Knotts et al., 2001), which we were able to confirm as the specific phosphorylation of S162 PR in response to progesterone was strongly decreased in the presence of CDK2 inhibition (Fig. 2H).

### 4. Functional Analysis

In order to investigate the dynamics of progestin signalling over time and with the aim of avoiding inherent biases generated from either technical approach, we combined the significantly regulated phosphorylation sites from both datasets (Fig. 1 and 2) resulting in a list of 420 unique phosphorylation sites within 390 proteins (Fig. S4A). PCA analysis of the samples reveals a clear separation of the phosphorylation data at 6 hours following hormone, given the majority of phosphorylation sites are rapid effect this separation of the latest time point may reveal changes in protein levels at this time point. The majority of these proteins showed the regulation of a single phosphorylation event with the exception of several highlighted proteins, including FAK, MAPT and EGFR (Fig. S4C). As in the individual analysis, phosphorylation sites were significantly regulated over several time points (Fig. S4D) and showed a switch from up-regulated sites early after hormone exposure to down-regulated sites at later time points (Fig. S4D). KEGG pathway analysis shows a significant enrichment in MAPK, PI3K-AKT, neurotrophin (TRK) and ERBB signalling pathways (Fig. 3A, Supplementary Table S2), in addition to pathways key in the progression of cancer, specifically cancer stem cells, such as focal adhesion (Fig. 3A).

**Fig. 3.**
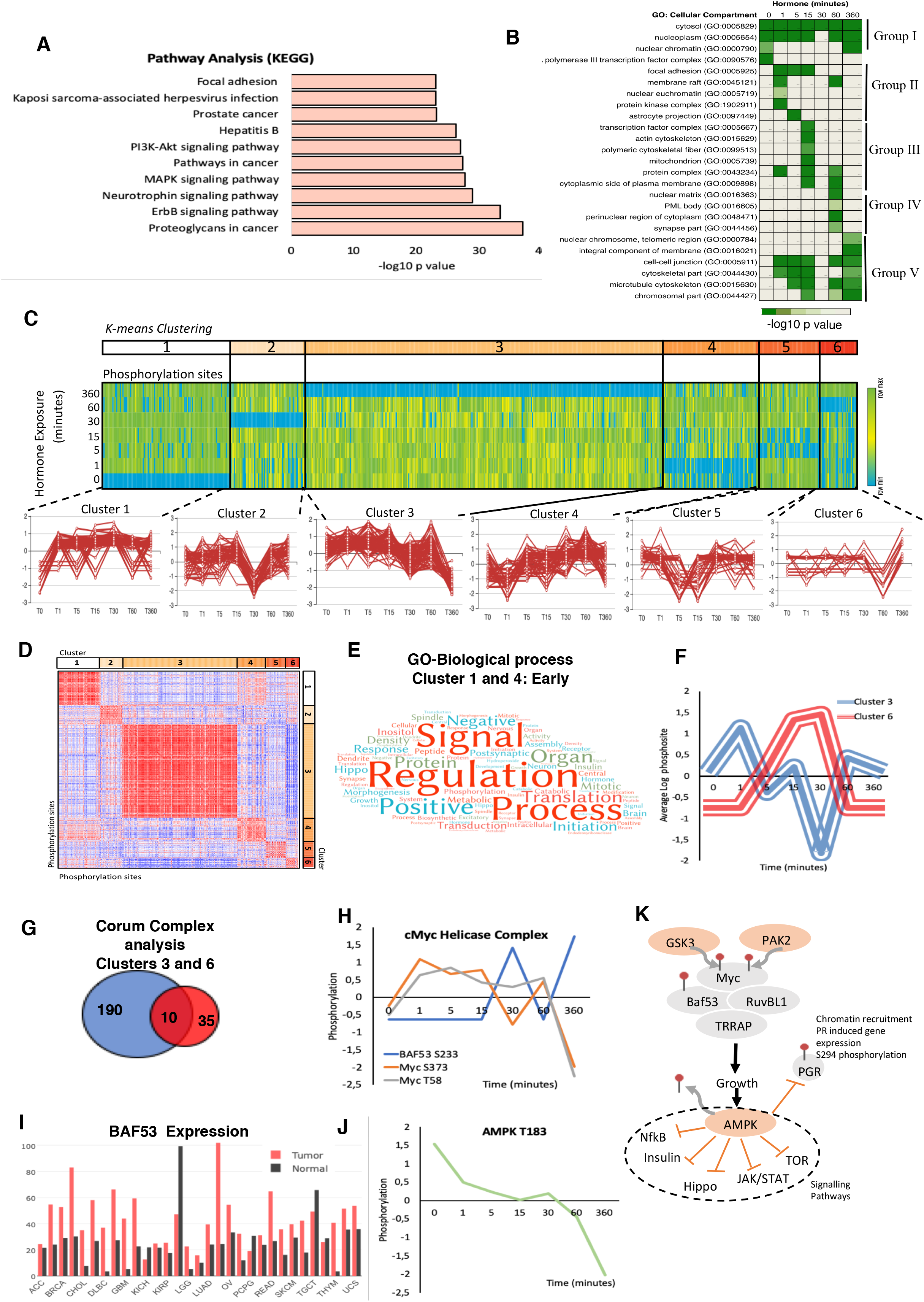
Combining Target Antibody Arrays and Shotgun Phosphoproteomic datasets following hormone. **A)** KEGG pathway enrichment analysis of proteins identified as regulated by phosphorylation in response to hormone. **B)** Cellular component analysis of phosphosites enriched per time point. Showing the hormone induced phosphorylation of the nucleoplasm and cytosol across all time points (group I) the activation of membrane raft proteins enriched at 1 minute (Group II) and phosphorylation of mitochondrial proteins enriched at 15 minutes (Group III), activation of nuclear structures; PML bodies and the nuclear matrix at 60 minutes (Group IV) and the activation of the cell-cell junctions and microtubules at 360 minutes (Group V). **C)** K mean clustering of all significantly regulated phosphorylation sites over time reveals 6 distinct clusters. **D)** Similarity matrix of clusters 1-6 reveals similar dynamics for clusters 1 and 4 and an opposing similarity in phosphorylation dynamics for clusters 3 and 6. Red indicates highly similar, well correlated, blue inversely correlated patterns of regulation. **E)** Word cloud showing the enrichment of GO-biological processes associated with proteins identified in similar clusters 1 and 4 “Early risers” which are regulated rapidly after hormone. **F)** Graph showing the opposing phosphorylation dynamic of proteins within clusters 3 and 6. **G)** Venn diagram showing the overlap of significantly identified Corum protein complexes identified in clusters 3 and 6. **H)** Phosphorylation dynamic in response to hormone of Myc S373, and T58 and BAF53 S233. **I)** Expression level of BAF53 in tumour versus normal tissue within the TGCA dataset. **J)** Phosphorylation of AMPL T183 decreases rapidly in response to hormone. **K)** Model showing the key role of AMPK dephosphorylation in response to hormone in breast cancer cells, AMPK dephosphorylation is required in order for subsequent signaling cascades including NFkB, insulin, Hippo, JAK/STAT and mTOR to continue and the phosphorylation of PR S294 to take place.

GO cellular component analysis reveals a dynamic pattern of specific cellular compartments over time (Fig. 3B, Supplementary Table S3). As expected, over the whole-time course, proteins are mainly found within the cytosol and nucleoplasm. However, prior to hormone exposure, phosphorylated proteins are enriched in RNA transcription repression complex and nuclear chromatin. The addition of hormone rapidly induces the phosphorylation of the membrane rafts, components of focal adhesion and protein kinases consistent with published works whereby signalling initiates from the plasma membrane. This transient phosphorylation of the membrane rafts diminish after 1 minute and is followed by the phosphorylation of transcription factors and proteins within the cytoskeleton (Fig. 3B). Interestingly, in line with our findings showing the generation of nuclear ATP synthesis independent of mitochondrial supplementation at 30 minutes after hormone we observe an enrichment in phosphorylated proteins located within the mitochondrial membrane at 15 minutes (Fig. 3B). The dynamic regulation of these proteins; CYB5B (cytochrome b5), the transcriptional activator ATF2 (Cyclic AMP-dependent transcription factor ATF-2), RPS6KB1 (Ribosomal protein S6 kinase beta-1) and PI4KB (Phosphatidylinositol 4-kinase beta) (Fig. S4F) may suggest an as yet undiscovered crosstalk between the nuclear and mitochondrial ATP synthesis pathways. Mitochondrial PR (PR-M) is a truncated isoform of the nuclear progesterone receptors PRB and PRA, which lacks the N-terminal DNA binding domain present in PRA and PRB but does contain the hinge region responsible for dimerization and the ligand binding domain (Price and Dai, 2015). PR-M has been shown to increase cellular respiration hence cell energy levels in response to ligand in various physiological situations and animal models (Dai et al., 2019). Therefore, the coordinated phosphorylation of proteins within the mitochondria in response to ligand (Fig. S4F) in breast cancer cells may provide an interesting insight into a possible crosstalk between mitochondrial PR-M and the nuclear receptors PRA and PRB.

At 60 minutes following hormone exposure the main localization of phosphorylation changes and shifts again to nuclear matrix proteins and proteins found within distinct regions of the nucleus, such as PML bodies (Fig. 3B group IV), which may be involved in the reorganization of chromatin in response to progestins (Le Dily et al., 2019). At 6 hours following hormone exposure cells enter the early stages of entering the cell cycle and movement is increased. This is also evident by the enrichment of phosphorylation sites in proteins within cell-cell junctions, the cytoskeleton and microtubules (Fig. 3B group V).

### 5. Protein class analysis

The majority of identified phosphorylated proteins (60%) were assigned to one class, however due to the promiscuous nature of enzymes nearly 40% were assigned to more than one class (Fig. S4G). Taking first only the parent class into account, we observed 5 distinct functions; 1) nucleic acid binding, 2) enzymes, 3) structural proteins, 4) protein modulators, and 5) proteins involved in signalling, membrane and cell-cell contacts (Fig. S4H). Each function class consists of sub-groups (Fig. S5A-F). The Nucleic Acid binding class includes DNA binding proteins, helicases, nucleases and RNA binding protein subgroups (Fig. S5A). The enzyme class is dominated by kinases but also includes histone modifying enzymes, hydrolases, ligases and oxidoreductases (Fig. S5D). The structural class is dominated by cytoskeleton proteins (Fig. S5F). The protein modulator class includes chaperones and various kinases and G proteins regulators (Fig. S5C). The cell signalling and the membrane/cell-cell contact classes are more complex and include many specialized proteins such as signalling molecules, receptors and transporters (Fig. S5B and E).

Gene Ontology of Biological Processes (GO-BP) and Molecular Function (GO-MF) showed an enrichment in signal transduction and general biological processes across the entire time course (Fig. S6A and B, Supplementary Tables S4 and S5). However, several interesting dynamic functions were identified. For instance, transcription co-factors, transcriptional repressors and transcription factor binding were already enriched 1 minute after hormone exposure (Fig. S6B) consistent with our previous observations of rapid transcription factor recruitment following hormone exposure (Nacht et al., 2016; Vicent et al., 2011). We observed enrichment in ATP binding and Adenyl-ribonucleotide binding after 5 and 60 min of hormone exposure (Fig. S6B), which may represent regulation of the two cycles of ATP dependent chromatin modifiers in response to progesterone (Vicent et al., 2009b; Wright et al., 2016). KEGG pathway analysis reveals a significant enrichment in signalling and in many cancer pathways, including Prostate, Glioma, CML, lung, AML, endometrial and pancreatic cancer, as well as focal adhesion and tight junctions (Fig. S6C). Annotated signalling cascades were significantly enriched at all time points in response to progestin, including MAPK, neurotrophin (TRK), ERBB, FC-receptor and insulin signalling (Fig. S6C).

### 6. Specific pathways: Roles of AMPK, insulin TNFa, and PIK3

K Means clustering analysis revealed six patterns of regulation over the time course (Fig. 3C). Similarity analysis of all phosphorylation sites within all clusters shows several interesting dynamics. First, “Early-risers” cluster 1 and 4, are positively correlated on the similarity matrix and show their initial increase in phosphorylation early at 1 and 5 minutes, respectively (Fig. 3D). GO-BP analysis of the proteins contained within these clusters shows an enrichment in signal regulation, and signalling cascades including Hippo, NFk-B and MAPK pathways (Fig. 3E, Supplementary Table S9). Second, clusters 3 and 6 show an opposing nature (negative correlation Fig. 3D). This antagonistic behavior of the two clusters is clearly shown averaging the signal of all phosphorylation within each cluster (Fig. 3F). Corum (comprehensive resource of mammalian protein complexes) analysis of the significantly enriched protein complexes contained within clusters 3 and 6 (Supplementary Table S6) showed that most protein complexes were enriched in one cluster or the other (Fig. 3G), likely representing crosstalk. Ten protein complexes were found to be enriched in both cluster 3 and 6, having phosphorylation sites within the same protein complex regulated in an opposite manner (Fig. 3G).

One such complex was the cMyc-ATPase-Helicase complex, which contains 5 proteins; cMyc, the chromatin remodeling component BAF53, the ATP-dependent helicases RUVBL1 and 2 (also known as TIP48 and 49) and the histone acetyltransferase, TRRAP. This complex is involved in chromatin organization, histone acetylation and transcriptional regulation (Park et al., 2002). Analysis of the phosphorylation sites showed that two sites (S373, T58) within Myc were increased early after hormone exposure, and decreased after 60 minutes, whereas one site of BAF53 (S233) shows the opposite dynamic (Fig. 3H). Database analysis also reveals a strong overexpression of BAF53 in tumour versus normal samples in multiple cancer types (Fig. 3I). Myc has an important role in breast cancer growth via the activation of AMPK (von Eyss et al., 2015).

The AMP-activated protein kinase (AMPK), exhibited a decrease in T183 phosphorylation in response to hormone. This site is phosphorylated by CAMKK1 or 2 (Hurley et al., 2005). AMPK is a master sensor, and its activation inhibits several kinase pathways including mTOR, NfkB, JAK/STAT, insulin and Hippo (Hadad et al., 2008; Montero et al., 2014; Yamaguchi and Taouk, 2020; Zhao et al., 2017). In addition, active AMPK inhibits the phosphorylation of PR S294, PR recruitment to chromatin and the activation of progesterone regulated genes (Wu et al., 2011). Activation of the kinase, specifically requires the phosphorylation of AMPK at T183 by CAMKKs, and de-phosphorylation of this site has been shown to be induced by estrogens and androgens in adipocytes (McInnes et al., 2012; McInnes et al., 2006). Previous published results and the data presented here suggests a model where AMPK must be silenced in order for regulatory pathways described and PR itself to be active, which is what we observe within 1 minute of progesterone stimulation (Fig. 3J and K).

Further in-depth analysis of the complexes which are regulated by phosphorylation in response to progestin revealed a full list of complexes with at least 2 proteins phosphorylated in response to hormone. One of them is the Sam68-p85 P13K-IRS-1-IR signalling complex, which encompasses the insulin receptor (INSR), the insulin receptor substrate 1 (IRS1), the KH domain containing transduction-associated protein 1 (Sam68) and the phosphatidylinositol 3-kinase regulatory subunit alpha (GRB1). This protein complex is involved in insulin signalling and has been proposed to provide a link between the PI3K pathway and other signalling cascades of insulin or p21/RAS (Sanchez-Margalet and Najib, 2001). We observed a dynamic phosphorylation of several sites within the complex (Fig. 4A), including 4 distinct phosphorylation events within IRS1, two of which peak at 1 minute (S312, S639) and two sites where the peak in phosphorylation is observed at 60 minutes (Y1179, and Y612). S312 has been shown to be directly phosphorylated by c-Jun N-terminal kinase (JNK1) in breast cancer signalling (Mamay et al., 2003) and this phosphorylation inhibits its interaction with IKKA. S639 is phosphorylated by mTOR has been linked to PI3K/Akt/mTOR signalling in breast cancer (Eto, 2010; Tzatsos, 2009) and effects the intracellular localization of IRS1 (Hiratani et al., 2005). Y1179 has been reported to be phosphorylated by IGF1R or INSR itself (Xu et al., 1995) and Y612 phosphorylation activates the interaction with PIK3R1 (Valverde et al., 2003).

**Fig. 4.**
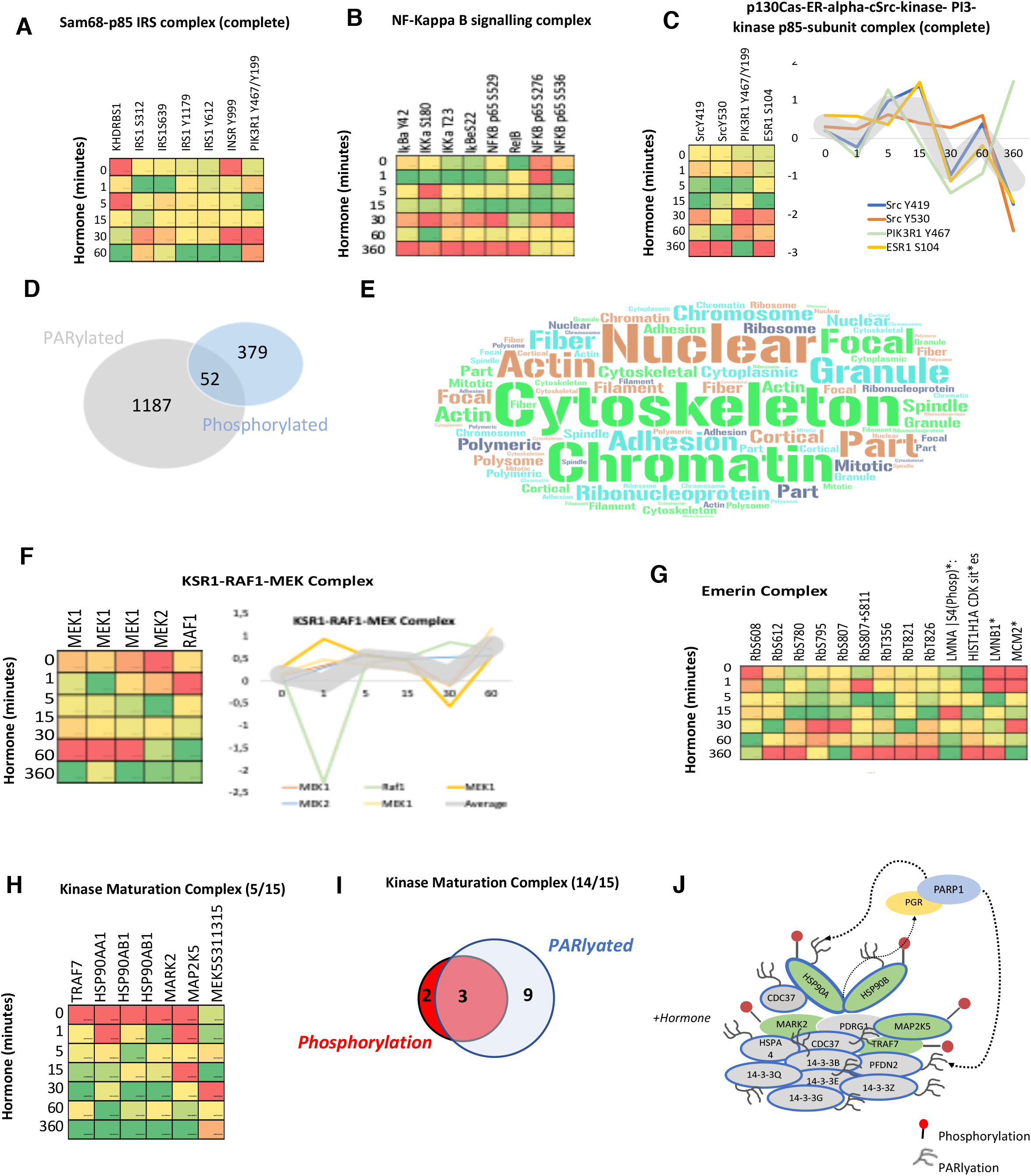
Protein Complex analysis and Overlap of PARylation and Phosphorylation in response to Progesterone. Heatmaps showing the phosphorylation of proteins within the Sam68-p85 IRS **(A)** and NF-kappa B **(B)** signalling complexes in response to hormone over time. **C)** Heatmap showing the phosphorylation of proteins of the p130 Cas-ER-Src-PI3K complex in response to hormone over time, the coordinated phosphorylation of each phosphosite individually is represented as a line graph (right panel). **D)** Venn diagram showing the overlap of proteins which contain either a phosphorylation site (379), PARylation site (1187) or both PTMs within the same protein after hormone exposure in breast cancer cells (52). **E)** Word cloud representation showing the GO-cellular component enrichment analysis of the 52 proteins identified as phosphorylated and PARylated in response to hormone (Fig. 4D). **F)** Heatmap showing the phosphorylation of components of the KRS1-RAF1-MEK signalling complex in response to hormone over time, all proteins shown are phosphorylated and PARylated and the dynamics of individual sites is shown on the right panel. **G)** Heatmap showing the phosphorylation proteins of the Emerin complex in response to hormone over time. **H)** Heatmap showing the phosphorylation of proteins (5/15) within the Kinase Maturation complex in response to hormone over time, phosphorylation dynamics of individual sites is shown (lower panel). **I)** Venn diagram showing the phosphorylation and or PARylation of proteins contained within the kinase maturation complex; 14/15 protein components of the complex contain at least one of the PTMs. **J)** Schematic representation of the complex components, PARylated proteins are indicated by blue circle, Phosphorylated by red (right panel).

The phosphorylation of several components of the TNFa/NFkB signalling complex were also identified (Fig. 4B). This complex is involved in I-kB kinase/NF-kB signalling in tumour progression. Indeed, complex components IKKα, RelB and p52 are associated with decreased cancer-specific survival in ERa-positive breast cancer (Paul et al., 2018). This may be linked to the cancer stem cell niche, which we showed recently was present in T47D cells grown in 3D cultures (Pickup et al., 2019). NFkB regulates self-renewal in breast cancer stem cell (BCSC) models and deletion of IKKα in mammary-gland epithelial cells affects progestin-driven breast cancer (Schramek et al., 2010; Shostak and Chariot, 2011). Indeed, the upstream activator RANK ligand (RANKL) and hence the RANK pathway promotes mammary tumor formation, (Gonzalez-Suarez et al., 2010), (Schramek et al., 2010). Another example is the P130Cas-ER-cSrc-PIK3 kinase complex (Fig. 4C). Which has been shown to induce transcriptional changes in response to estrogen and mammary proliferation in breast cancer. The authors showed that estradiol triggers the association of ERa, c-Src, the p85 subunit of PI 3-kinase (PI3K) and p130Cas in a macromolecular complex and activates the c-Src kinase leading to p130Cas-dependent Erk1/2 phosphorylation (Cabodi et al., 2004; Cabodi et al., 2006). Given the similarity of the phosphorylation dynamics, peaking early at 5 and 15 minutes across Src, PIK3 and ESR1 within the complex induced by progestin (Fig. 4C right panel) this may (similarly to the induction by estrogen shown by others) present a novel ERa-ERK-cSrc activation mechanism in response to progestin in breast cancer cells.

### 7. Crosstalk between Progestin induced Phosphorylation and PARylation

Recently, it has been shown that the majority of PARylation events on eukaryotic nuclear proteins take place on serine residues rather than acidic residues as previously accepted (Bonfiglio et al., 2017a; Bonfiglio et al., 2017b; Leidecker et al., 2016; Liu et al., 2017; Martello et al., 2016; Messner et al., 2010). Given the enrichment of serine in the phosphorylation dataset (Fig. 2D) and the importance of both PTMs in progestin gene regulation (Wright et al., 2016), we investigated the overlap between PARylation sites and phosphorylation sites within protein complexes. We identified 52 proteins, which were both phosphorylated and PARylated in response to progestins in breast cancer cells (Fig. 4D). Cellular component analysis of this set of 52 proteins indicates a significant enrichment in nuclear, cytoskeleton and chromatin contained proteins (Fig. 4E, Supplementary Table S7), in line with the well described role of PAR in the nucleus, nuclear organization, chromatin organization, relaxation and transcriptional regulation (Hassa et al., 2006; Hoch and Polo, 2019; Leung, 2014; Thomas and Tulin, 2013). This finding may indicate a crosstalk between PARylation and phosphorylation with regards to nuclear structure and chromatin organization. Analysis of complexes significantly enriched within this group of proteins revealed 12 protein complexes (Supplementary Table S8), which contained proteins both PARylated and phosphorylated. One of them is the KSR1-RAF1-MEK complex composed of MEK1 and 2, both PARylated and phosphorylated, and RAF1 which is phosphorylated (Fig. 4F). This complex is involved in the MAKPKKK cascade, and in response to EGF it activates BRAF mediated phosphorylation of MEK1, at 3 sites, and MEK2, which activate MAPK1 and 3 (McKay et al., 2009). In our dataset we observe a clear change in phosphorylation of all members of the complex in response to progestin (Fig. 4F right panel).

We also observed the phosphorylation and PARylation of the Emerin complex (Fig. 4G). This complex is involved in DNA replication, transcription and structural integrity of the nucleus, specifically of the inner nuclear membrane (Holaska and Wilson, 2007). Depletion of Emerin results in changes in the organisation and dynamics of the nucleus, increased chromatin mobility and a mis-localisation of chromosome territories (Ranade et al., 2019). Within this complex we find proteins phosphorylated, PARylated, or phosphorylated and PARylated (Fig. 4G). Given the role of PAR in the structure of the nucleus, this complex may present and interesting example for studying the PARylation, phosphorylation crosstalk.

As discussed, prior to hormone exposure PR is present in an inactive complex with the HSP70 and 90 proteins as part of the Kinase Maturation Complex. We know that progestins promote the phosphorylation and dimerization of the receptor and we found that phosphorylation of the HSP90 and 70, along with other members of the complex, is initiated within 1 minute of hormone exposure (Fig. 4H), again showing a rapid and concerted phosphorylation of several members of the complex (Fig. 4H). In addition, the HSPs are also PARylated as compared to other components where only phosphorylation (MARK2, MAP2K5) or PARylation (14-3-3 components) are present (Fig. 4I and J). Further investigation regarding the crosstalk between PARylation and phosphorylation within protein complexes will be the focus of future studies.

### 8. Progesterone Signalling Network Generation

In order to understand the crosstalk between the signalling pathways activated by Progesterone in breast cancer cells, a Protein-Protein Interaction (PPI) network was generated using all identified phosphorylation sites (Supp. File: Network session 2), based on known PPI (evidence based). The resulting network consists of 427 nodes (proteins) and 4309 unique interactions (edges) (Fig S7A). Pathway analysis was carried out on this network, using Genemania™, and 23 statistically significant (p<0.01) pathways were identified (Supplementary Table S9 and S10). The proteins and interactions (nodes and edges) associated with each pathway were selected and new networks generated (Supp. File: Network session 2). Several of which are shown in Fig. S7B-G and discussed briefly below.

One such pathway, the Fc receptor signalling pathway was identified as enriched (Fig. S6C) and the PPI network is shown in Fig. S7B. Fc receptors are cell surface proteins that recognize the FC fragment of antibodies, mainly on immune cells. However, recent studies have shown that different subsets of Fc receptors may play a role in tumour cells (Nelson et al., 2001). In particular, it was shown that T47D cells express the FcyRI (CD64) These FC-receptor expressing breast cancer cells can activate the tyrosine kinase signal transduction pathway. Indeed, T47D cells treated with selective tyrosine kinase inhibitors do not proliferate in a FC receptor-tyrosine kinase signalling dependent manner (Nelson et al., 2001).

As mentioned before (Fig. S6C), another pathway identified as activated in response to progestin is the ERBB-EGF network (Fig. S7D and H). ERBB2 (HER2) is overexpressed in 15-20% of breast cancer in response to EGF activation, and plays a major role in EMT (Elizalde et al., 2016). PR interacts with ERBBs and induces the translocation of ERBB2-PR-STAT3 complex to the nucleus. ERBB2 acts as a co-activator of STAT3 and drives the activation of progestin regulated genes, especially genes such as Cyclin D1 that do not contain a HREs (Beguelin et al., 2010; Hsu and Hung, 2016). Blocking PR signalling in PR-ERBB2 positive breast cancer patients has been suggested as a treatment (Proietti et al., 2009). The ERBB pathway may represent a new mechanism for further study to understand the activation of these “non-classical” PR dependent genes in response to progestin. In addition to the role of ERBB2, the role of ER activation in response to progesterone in breast cancer cells is also critical, as shown in Fig. 4C we observe the coordinated activation of the ESR1-Src-PIK3 complex peaking at 15 minutes following hormone exposure. The phosphorylation site of ESR1 which increases is S104. ER S104 phosphorylation is essential for ER activity (Thomas et al 2008) and it has been suggested that hyperphosphorylation of ER at these sites may contribute to resistance to tamoxifen in hormone receptor positive breast cancer (Leeuw et al 2011, Jeffreys et al 2020, Skliris et al 2010). ER S104 phosphorylation by ERK has been shown previously in response to estrogen and EGF but not progesterone exposure. In addition, ER S104 has been implicated in mTOR signaling (Alayev et al 2016). Given the phosphorylation of ER, the dynamic activation of the ER membrane complex and the role of mTOR in AMPK and insulin signalling described earlier (Fig. 3J and 4C) this phosphorylation site may present a key step in the cellular response to progesterone in breast cancer.

A pathway exhibiting strong activation by progestins is the Insulin signalling (Fig. S6C, Fig. S7F). Insulin-like growth factors (IGFs) and progestins both play a major role in normal mammary gland development and R5020 has been shown to induce the expression of insulin receptor substrate-2 in MCF7 cells (Cui et al., 2003a; Cui et al., 2003b). Moreover, IGF signalling via IRS2 is known to be essential for breast cancer cell migration. It has also been shown that R5020 pretreatment followed by IGF stimulation increases binding of IRS to PI3K-p85 regulatory complex, which in turn activates ERK and AKT signalling (Ibrahim et al., 2008). Interestingly, not only do we observe the activation of the insulin pathway in network analysis (Fig. S7F, but the coordinated phosphorylation of all members of the IRS-PIK3 complex was also identified (Fig. 4A), indicating that indeed progesterone stimulation of breast cancer cells activates the not only the insulin pathway but the coordinated regulation of complexes within it.

## Discussion

The data presented in this paper provides a source of knowledge for the scientific community with regards to progesterone induced gene expression, and the signalling pathways involved. We have shown the rapid induction of phosphorylation using two distinct technologies (Fig. 1 and 2). Pathway analysis showed a strong enrichment in pathways associated with cancer, known and novel Pg-dependent signalling events (Fig. 3A). But also identified signalling pathways not previously known to mediate progesterone action in breast cancer cells, such as MNK1/EIF4E pathway and the connection between CDK2, Cdc25 and the MAPK pathway.

Cellular component analysis confirmed our expectations and the statistically significant activation of the cell membrane within 1 minute of hormone exposure (Fig. 3B), but also revealed a consistent (over all members) phosphorylation peak within proteins associated within the mitochondria at 15 minutes after hormone exposure (Fig. 3B and S4E). Mitochondrial activation in response to progesterone in breast cancer cells has not been extensively studied yet. Indeed ATP synthesis 45-60 minutes after hormone stimulation is independent of mitochondrial involvement (Wright et al., 2016). However, there are some interesting findings in the literature. Following the observation that the PR negative cell line MCF10A exhibits a progestin-induced cell proliferation (Kramer et al., 2006). Behera and colleagues showed that MCF10A respond to R5020 with and increase in mitochondrial activation (Behera et al., 2009). Given the absence of the nuclear PR in these cells they hypothesized that the activation of progestin-induced cell growth was due to non-genomic metabolic effects, mediated by a yet undiscovered receptor. We propose that the observed mitochondrial activation in T47D in response to progestin (Fig. 3B and S4E) suggests the existence of a third and interconnected hormonal signal transduction pathway via the mitochondria (Demonacos et al., 1996; Hatzoglou and Sekeris, 1997).

Dynamic phosphorylation analysis over time reveals distinct groups or clusters of phosphorylation events which follow a similar time response; such as early risers, sustained or late (Fig. 3C). Similarity analysis of these dynamic phosphorylation sites reveals some interesting crosstalk between protein complexes not previously identified as players in progesterone signalling in breast cancer cells (Supplementary Table S6), and complexes where a mobilization of phosphorylation (showing similar dynamics) was observed within the whole macromolecular complex; such as PIK3, NFkB (Fig. 4B and C).

Overlap of phosphorylation sites with existing PARylation, revealed 52 proteins for which both phosphorylation and PARylation was found. The data also clearly shows an enrichment in protein complexes that play a role in the structural organisation of the nucleus (Fig. 4D and E), specifically the Emerin complex and Lamin (Fig. 4G). The key location of these complexes at the nuclear membrane, suggests that perhaps these two PTMs may affect and play a role in the dynamic structure of the nucleus. This could be explored in the future by global chromatin proximity Hi-C experiments. The complexes identified in this study and the dual post translational modification of proteins with known important roles within the cell may provide exciting opportunities for future studies which aim to understand the crosstalk between Serine PARylation and phosphorylation in the context of nuclear architecture, signalling and breast cancer progression (Supplementary Table S7).

In addition to pathway analysis at the single network level (Fig. S7), wherein we identified pathways such as insulin, Fc-receptor and ERBB signalling, it is also clearly important to consider the connection between pathways and networks as a whole. One such example, is the connections between the phosphorylation events within the cytoskeleton, membrane raft and proteins associated with cell adhesion. We observe phosphorylation events in multiple proteins within both cell adhesion and the membrane raft, forming tight strongly connected PPI networks (Fig. 5A). A network merge of these two pathways reveals 4 key proteins which are present in both (JAK2, SRC1, LYN and KDR), indicating a strongly connected network (Fig. 5B), which in addition to common members exhibits a large first neighbour selection between the two initial pathways (selection of only direct PPI) (Fig. 5B right panel) with a similar phosphokinetic pattern (Fig. 5C). Incorporation of the significantly enriched phosphorylated proteins within the cytoskeleton (Fig. 3B, Fig. S6D, Supplementary table S3) into the merged network (Fig. 5B) results in a larger global connected network (Fig. 5D) which supports the activation within the membrane proteins after 1-minute following hormone exposure that triggers the subsequent cascades of phosphorylation in the cytoskeleton (Fig. 5E). These findings clearly show the importance of studying the pathways not in isolation, but rather in connection with each other.

**Fig. 5.**
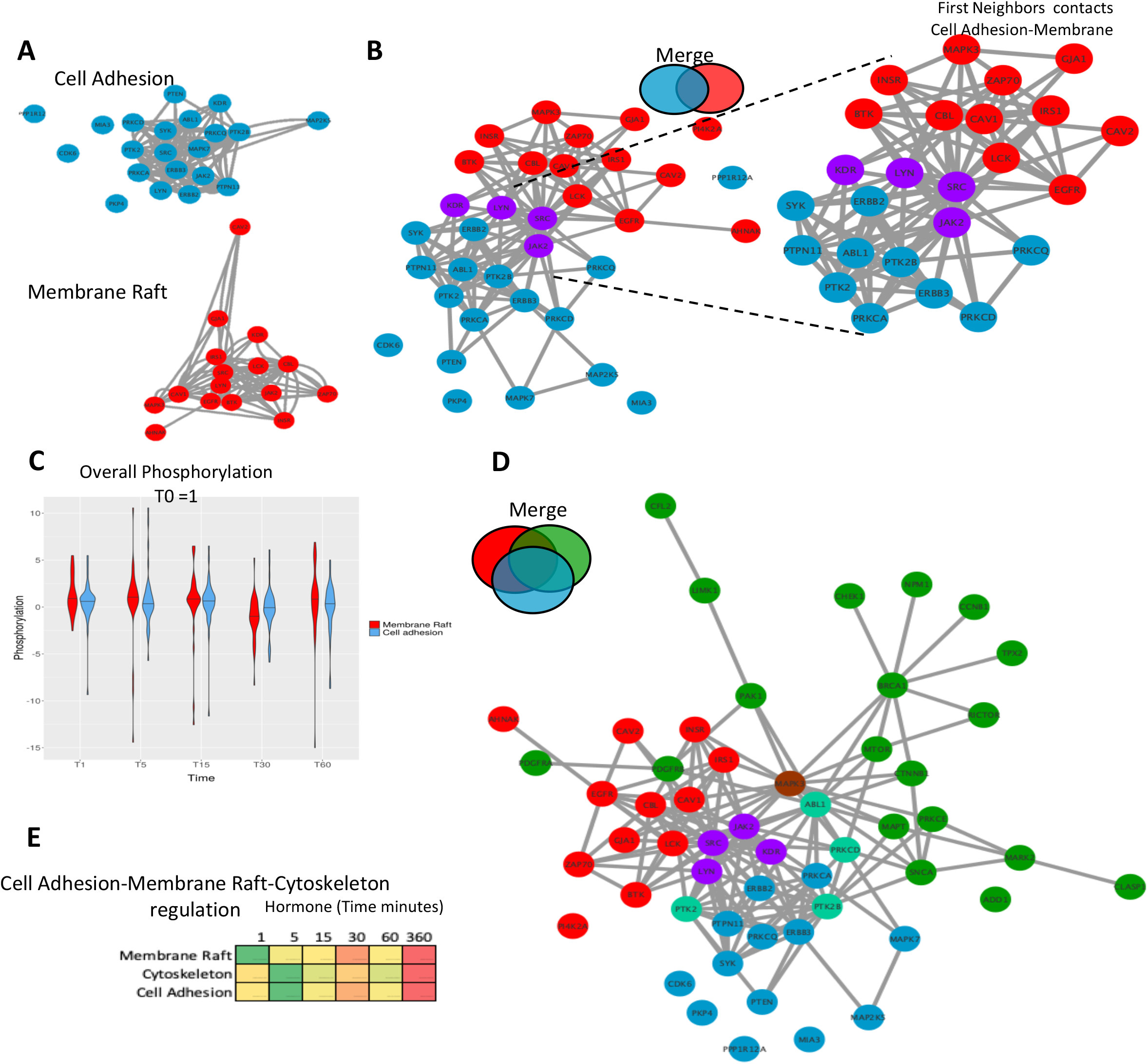
Combining PPI networks from distinct cellular compartments reveals a coordinated crosstalk. **A)** PPI network showing the significantly regulated phosphorylated proteins located in cell adhesion (blue) and the membrane raft (red) identified in response to hormone. **B)** Merge of the cell adhesion network (Fig. 5A, blue) and membrane raft network (Fig. 5A red). The two networks connect based on known PPI however no protein was identified as annotated in both sets. This integration of the two networks is highlighted (right panel) where proteins from each network were selected based on having a first neighbour with a protein of the other network. **C)** Violin plot showing the average phosphorylation of proteins over time in response to hormone within the membrane raft or cell adhesion networks. Data is normalised to time 0=1. **D)** Merge of Cell Adhesion-Membrane (Fig. 6B) and the cytoskeleton networks. The two networks are merged based on known PPI. Proteins annotated in more than one function are coloured based on the Venn diagram (i.e. cytoskeleton and cell Adhesion; light green, membrane raft and cytoskeleton; brown). **E)** Heatmap showing the average phosphorylation of all proteins within each network in response to hormone over time, showing the activation of the membrane raft first at 1 minute followed by the cytoskeleton and cell adhesion.

Other examples of connectivity are observed between the ERK subgroup and the MAPK cascades and the FC Receptor and TRK signalling. ERK and MAPK form a strong network (Fig. 6A and B). As discussed, earlier Fc-receptor signalling shows a strong activation (Fig. S7B). The tropomyosin receptor tyrosine kinases (TRKs) are primarily known for their roles in neuronal differentiation and survival. However, increasing evidence shows that TRK receptors can be found in a host of mammalian cell types to drive several cellular responses (Huang and Reichardt, 2003; Reichardt, 2006) Aberrations in TRK signalling, which can occur through events such as protein overexpression, alternative splicing, or gene amplification, can lead to disease such as cancer (Jin, 2020; Meng et al., 2019; Regua et al., 2019). The receptor tyrosine kinase NTRK2, activates GRB2-Ras-MAPK cascade in neurons and increases secretion by epithelial cells in culture in response to estrogen or progestin treatment and NTRK2 was identified as differentially expressed between stromal and epithelial breast cells which may have implications in invasion and metastasis (Wang et al., 2020) Merging of Fc-receptor and TRK signalling pathways (Fig. 6C) shows a strong protein overlap and a dense connected network with FOXO1 at the center (Fig. 6D). and FOXO1 is phosphorylated after 1 to 5 minutes of progestin exposure (Fig. 6E). Phosphorylation of FOXO1 by PKB/Akt has been shown to be important for the binding to 14-3-3 proteins on chromatin (Dobson et al., 2011; Pennington et al., 2018; Tzivion et al., 2011; Tzivion and Hay, 2011). The role of these two pathways in progesterone induced gene regulation has not been shown previously. FOXO factors have a key role to play in tumour resistance to therapy and patient outcome (Bullock, 2016). Interestingly from a clinical perspective stratifying patients based on either the expression levels of NTRK2 (TRKB) or FOXO1 is predictive of a good prognosis (overall survival) in breast cancer, similar to prognosis based on PR expression (Fig. 6F and G).

**Fig. 6.**
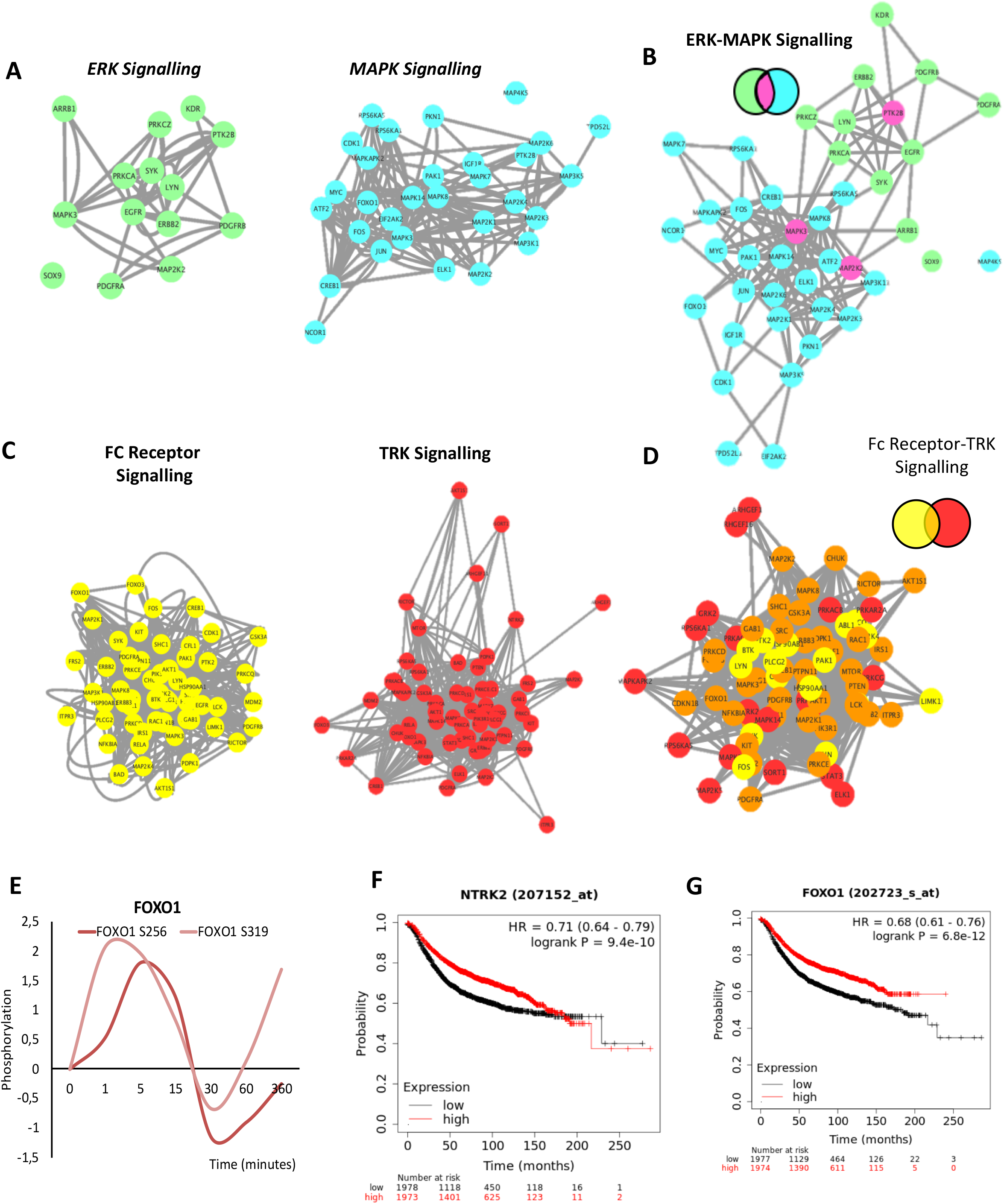
Network Integration of signalling networks identified in response to hormone. **A)** PPI network showing the phosphorylated proteins present within the ERK signalling cascade (green) and the MAPK cascade (blue) identified in response to hormone. **B)** Merge of ERK-MAPK networks (Fig. 6A). The two networks are merged based on known PPI. Proteins annotated in both pathways are coloured based on the Venn diagram (fuchsia). **C)** PPI network showing the phosphorylated proteins present within the FC-receptor (yellow) and TRK-neurorophin (red) signalling pathways (yellow) identified in response to hormone (left and middle panel). **D)** Merge of FC-receptor and TRK neurotrophin networks. The two networks are merged based on known PPI. Proteins annotated in both pathways are coloured based on the Venn diagram (orange). **E)** Rapid and coordinated phosphorylation of FOXO1 S256 and FOXO S319 in response to hormone. Kaplan Meyer overall survival of patients stratified based on the expression of NTRK2 **(F)** and FOXO1 **(G)** in breast cancer patients (p=9.4E-10 and 6.8e-12 respectively). All networks, PPIs and integrated cascades are supplied in Cytoscape session 2.

## Conclusions

The examples described here in addition to other examples contained within the data for future discovery show the importance of network connectivity in trying to understand not only individual proteins or pathways but the significant overlap between the pathways within the signalling network activated by progesterone in breast cancer. Further analysis of the detected connections and identification of the key regulators may provide a source of targets for drug discovery aiming at the treatment of hormone receptor positive breast cancer patients.

## List of abbreviations

ADPR: ADP ribose
AKT: RAC-alpha serine/threonine-protein kinase
AMPK: AMP activated protein kinase
ARTD1: ADP-ribosyl transferase 1
ATF: Cyclic AMP-dependent transcription factor
ATP: Adenosine triphosphate
BAF: BRG1-associated factor
BRAF: Serine/threonine protein kinase B-raf
CAMKK: Calcium/calmodulin-dependent protein kinase kinase
CARM1: Coactivator-associated arginine methyltransferase 1
cAMP: cyclic AMP
CDK: Cyclin dependent kinase
CSC: Cancer stem cells
CYB5B: Cytochrome b5 type B
DMBA: 7, 12-Dimethylbenz[a]anthracene
EGFR: Epidermal growth factor receptor
EMT: Epithelial to mesenchyme transition
ERK: Extracellular signal-regulated kinase
ERBB: Receptor tyrosine-protein kinase
ESR1/ER: Estrogen Receptor
FAK: Focal adhesion kinase
GRB: Growth factor receptor-bound protein
GSK3: Glycogen synthase kinase-3
H1: Histone H1
HRE: Hormone Responsive Elements
HSP: Heat shock protein
IGF: Insulin-like growth factors
IKKa: Inhibitor of nuclear factor kappa-B kinase
INSR1: Insulin receptor
IRS2: Insulin receptor substrate 2
JAK: Janus Kinase
KDR: Kinase insert domain receptor
KIP: Kinase inhibitor protein
KSR1: Kinase suppressor of Ras1
LYN: Lck/Yes-related novel protein tyrosine kinase
MAPK: Mitogen-activated protein kinase 1
MAPT: Microtubule associated protein Tau
MEK: MAPK/ERK kinase 1
MLL: Myeloid/lymphoid or mixed-lineage leukemia
MNK1: MAPK signal-integrating kinase 1
MPA: Medroxyprogesterone acetate
MSK1: Mitogen- and stress-activated protein kinase
MTOR: Mammalian target of rapamycin
NAD+: nicotinamide-adenine dinucleotide
NFkB: Nuclear factor NF-kappa-B
NUDT5: Nucleoside diphosphate-linked moiety X motif 5
NURF: Bromodomain and PHD finger-containing transcription factor
Pg: Progesterone
PARP1: Poly-ADP-ribose polymerase
PARylation: Poly-ADP-Ribosylation
PARG: Poly (ADP-ribose) glycohydrolase
PCA: Bromodomain adjacent to zinc finger domain protein 2B
PI4K: Phosphatidylinositol 4 kinase
PKN: Prior knowledge network
PLK: Serine/threonine-protein kinase
PML: Promyelocytic leukemia protein
PR/PGR: Progesterone Receptor
PPI: Protein-protein interactions
PRE: Progesterone responsive element
PTK: Protein-tyrosine kinase
PTM: Post translational modification
RAF: RAF protooncogene serine/threonine protein kinase
RANK: Receptor activator of NF-KB
RB1: Retinoblastoma-associated protein
RPS6KA1: Ribosomal protein S6 kinase alpha-1
RAS: GTPase HRas
RUVBL1: Nuclear matrix protein 238
Sam68: KH domain containing transduction associated protein 1
SRC: Proto-oncogene tyrosine-protein kinase Src
SRRM: Serine/arginine repetitive matric protein
STAT: Signal transducer and activator of transcription
TIP48/89: TIP60-associated protein 48/49
TNF: Tumour necrosis factor
TP53: Tumor suppressor p53
TRK: Tyrosine kinase receptor
VEGFR: Vascular endothelial growth factor receptor

## Declarations and Author contributions

The authors declare no competing interests and concept for this manuscript to be published. This work was supported by XXX. The authors would like to acknowledge all members of the core facilities Proteomic and Genomic Units in the CRG. Experimental design; R.H.G.W and M.B. Bioinformatic Analysis; J. Q. O., J.C.C and R.W. Manuscript writing and editing; R.H.G.W and M.B. Experiments; R.W.

## Supplementary Figure Legends

**Fig. S1. Prior Knowledge Network (PKN) Progesterone Signalling**.

**A)** Edge directed PKN network was manually curated from the literature. Annotated phosphorylation events, interactions, dissociations and cellular compartment are indicated. Network is available as a cytoscape network session (cys) or interaction (sys) file containing references for all edges present as shown in B) (See Supplementary File Network 1).

**Fig. S2. Antibody Array controls and data analysis**.

**A)** Schematic indicating the experimental procedure, quality control checks, and filtering applied to the antibody array experiments. **B)** Number of phosphosites identified per protein. Tau, PTK2, RPS6KA1 and RB1 are highlighted as they have multiple sites identified**. C)** Functional classification of the proteins identified as significantly phosphorylated at each time point **D)** KEGG pathway analysis, showing significant pathways (-log_10_ p-value) at each time point. **E)** Heatmap representation of GO biological process data, showing significant (-log_10_ p-value) processes at each time point. **F)** Heatmap representation of GO molecular function data, showing significant (-log_10_ p-value) functions at each time point.

**Fig. S3. Phosphoproteomic data acquisition and controls**.

**A)** Correlation of triplicate samples from each of the time points. **B)** Number of phosphosites identified per protein, the proteins showing multiple sites per protein are highlighted. **C)** KEGG pathway analysis, showing significant pathways (-log_10_ p-value) at each time point. **D)** Heatmap representation of GO-biological process data, showing significant (-log_10_ p-value) processes at each time point. **E)** Heatmap representation of GO-molecular function data, showing significant (-log_10_ p-value) functions per time point.

**Fig S4. Combining Antibody Array and Phosphoproteomic LC-MS-MS datasets**.

**A)** Schematic representation showing the methodology and overlap combining antibody array and LC-MS-MS datasets. **B)** PCA analysis of phosphorylation datasets. **C)** Number of phosphosites identified per protein, the names of proteins showing multiple sites per protein are highlighted. **D)** Venn diagram showing the overlap of phosphosites per time point. **E)** Up and down regulated phosphorylation sites identified per time point. **F)** Phosphorylation levels of the proteins identified as significantly regulated after hormone located within the mitochondria. **G)** Analysis of the number of functions to which each unique protein was assigned **H)** Venn diagram showing the overlap of protein functional class; Enzymes, Structural protein, Membrane-cell-cell contact, protein modulators and proteins with nucleic acid binding capacities.

**Fig S5. Functional Analysis of Proteins Identified.**

All proteins were assigned one or more function based on GSEA database. Both Parent (outside/title), and children (within) are shown for each class and the proteins identified within that sub-group are shown. Nucleic acid binding **(A),** Membrane/Cell-cell contact **(B),** Protein Modulators **(C),** Enzymes **(D),** Cell signalling **(E),** and Structural proteins **(F)**.

**Fig S6. Gene Ontology and Pathway analysis of combined dataset.**

**A)** Heatmap representation of GO biological process data, showing significant (-log_10_ p-value) biological processes enrichment based on the protein phosphorylation at each time point. **B)** Heatmap representation of GO molecular function enrichment, showing significant (-log_10_ p-value) functions at each time point following hormone. **C)** KEGG pathway analysis, showing significant pathways (-log_10_ p-value) enriched at each time point following hormone exposure. **D)** Protein protein interaction (PPI) network generated using proteins identified as phosphorylated following hormone and were assigned as cytoskeleton located.

**Fig S7. Pathway Network Generation in Breast Cancer cells in response to Hormone**.

**A)** Protein protein interaction (PPI) network was generated using full phosphorylation dataset encompassing 321 proteins (Supplementary Material Network session 2) in Cytoscape using Genemania™ only considering protein-protein interactions with experimental evidence (Supp. Materials and methods), each node represents and individual protein and interactions are represented by edges. Functional analysis was carried out to identify key pathways enriched within the full network (Full list Supplementary Table 16). Individual networks were generated from each function individually and are available within additional Network session 2. Graphs of several pathways determined to be enriched within the dataset are shown **B)** Fc receptor, **C)** MAPK, **D)** EGF **E)** ERK **F)** Insulin, **G)** TRK signalling, **H)** ERBB.

## Supplementary Table Legends

**Supplementary Table S1**

Uniprot IDs of phosphorylated proteins identified in response to hormone. Time after hormone (minutes), data is normalized 0-1 row maximum and minimum.

**Supplementary Table S2**

KEGG pathway enrichment; the pathway term, p value and the proteins associated with the pathway are shown.

**Supplementary Table S3**

Cellular component enrichment analysis of phosphorylated proteins. The time after hormone in which they peak, the adjusted p value, and proteins associated with each specific cellular component are given.

**Supplementary Table S4**

Gene Ontology Biological Process enrichment analysis of phosphorylated proteins. The cluster in which the term is enriched, the adjusted p value, and proteins associated with each specific biological process are given.

**Supplementary Table S5**

Gene Ontology Molecular Function enrichment analysis of phosphorylated proteins. The cluster in which the term is enriched, the adjusted p value, and proteins associated with each specific molecular function are given.

**Supplementary Table S6**

Corum enrichment analysis of phosphorylated proteins. The p-value, and proteins associated with each complex are given.

**Supplementary Table S7**

Cellular component enrichment analysis of phosphorylated and PARylated proteins. The adjusted p value, and proteins associated with each specific cellular component are given.

**Supplementary Table S8**

Corum enrichment analysis of phosphorylated and PARylated proteins. The p-value, and proteins associated with each complex are given, phosphorylated proteins are highlighted in yellow.

**Supplementary Table S9**

Genemania analysis of phosphorylated proteins, all protein IDs are listed along with the GO: IDs for which they are associated.

**Supplementary Table S10**

Genemania analysis of phosphorylated proteins, the pathways enriched in Network 2 are shown. The q value and the number of occurrences in the network versus the occurrences in the Network are shown.

## Additional Files

### Network Session 1

Edge directed PKN network was manually curated from the literature. Annotated phosphorylation events, interactions, dissociations and cellular compartment are indicated.

### Network Session 2

Protein protein interaction (PPI) network was generated using full phosphorylation dataset encompassing 321 proteins in Cytoscape using Genemania™ only considering protein-protein interactions with experimental evidence each node represents and individual protein and interactions are represented by edges. Functional analysis was carried out to identify key pathways enriched within the full network. Individual networks were generated from each function individually and are available as unique networks within Network session.

## Methods

### Cell culture

The hormone receptor positive breast cancer cell line T47D^M^ (REF) was used in all experiments unless otherwise stated. T47D^M^ cells were routinely grown in RPMI (Supplemented with 10% fetal bovine serum (FBS), penicillin/streptomycin (pen/strep), L-glutamine (L-glut) as previously described (Wright et al., 2016). For hormone induction experiments, cells were seeded at a concentration of 5 × 10^6^ per 150mm cell culture dish in RPM1 white (15% charcoal stripped FBS, Pen/strep, L-glut) for 48 hours. 16 hours prior to hormone induction (10nM R5020), medium was replaced with RPM1 white (0% FBS, Pen/strep, L-glut). Samples were harvested at the time points indicated.

### BCA Assay

The total protein content of the samples was calculated prior to antibody array, mass spec or western blotting analysis using BCA assay (Thermo Fisher, catalogue number 23227) according to manufactures instructions.

### Protein Visualisation

Changes in phosphorylation sites within individual proteins identified was confirmed by western blotting as previously described (Nacht et al., 2016) using specific antibodies; phospho S162 PGR (Abcam ab58564), and as a loading control, total PGR (Santa Cruz H190), total CDK2 (Santa Cruz sc-163) or CDK2 phospho T160 (Abcam ab47330).

### Antibody Microarray

Phosphorylation antibody array analysis was carried out by Kinexus™ using Kinexus ™ Antibody Microarray (KAM) technology. For each replicate, 50ug of protein lysate was prepared and samples prepared by Kinexus™ in house. Signal quantification was performed using ImaGene 8.0 from BioDiscovery (El Segundo, CA, USA). Background corrected raw intensity data was logarithmically transformed with base 2 and Z scores calculated (Cheadle et al., 2003). Any poor-quality spots based on morphology and background, were flagged as unreliable and removed from any subsequent analysis.

### Mass Spec Sample Preparation

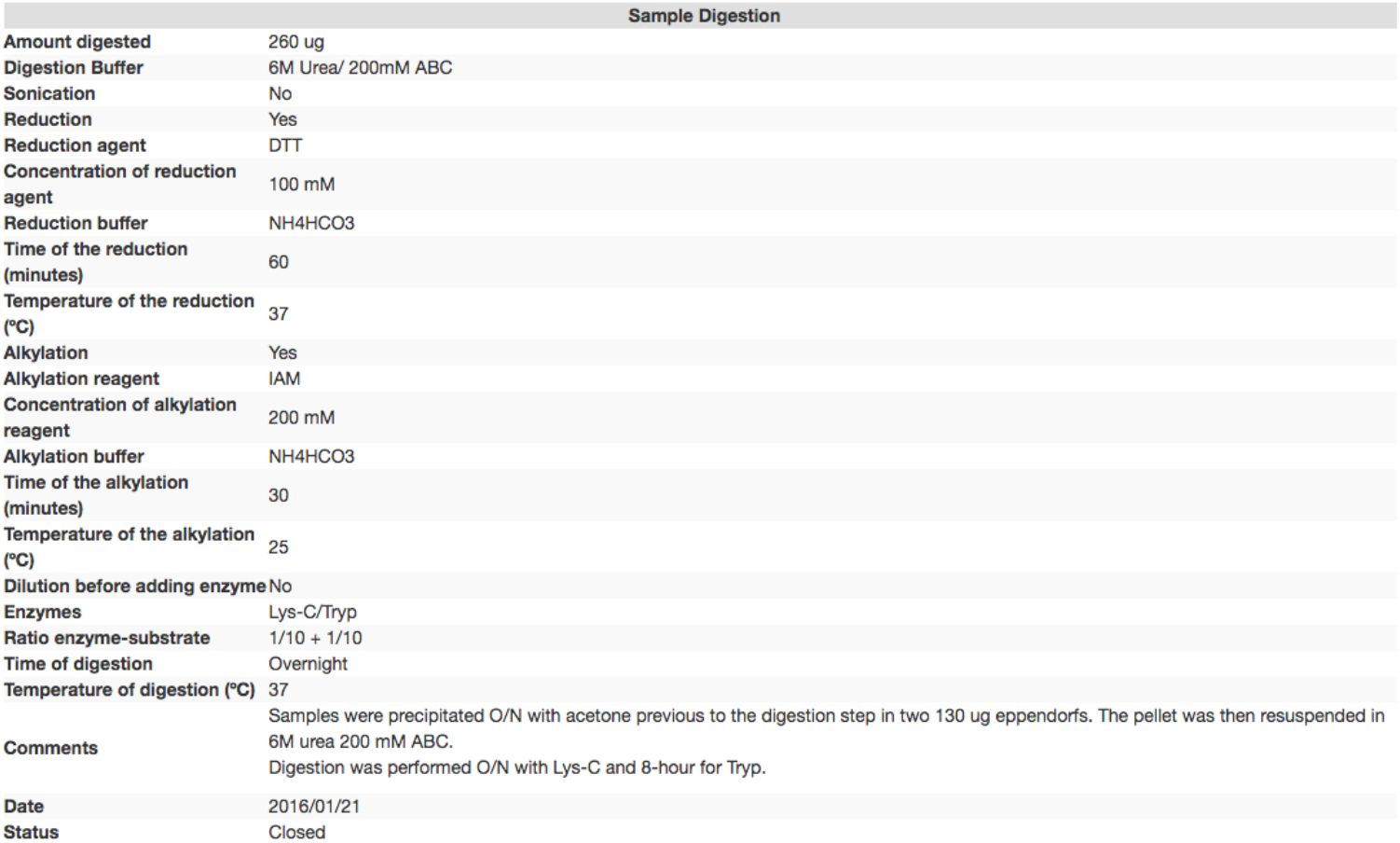

### Bioinformatic Procedures

Gene Ontology (GO) GO-Biological process (GO-BP), GO-Molecular Process (GO-MF), KEGG and Biocarta pathway analysis of networks was carried out using GeneMania application (Warde-Farley et al., 2010) within Cytoscape. Clustering analysis, similarity analysis was carried out using GeneE. Network analysis was performed using Cytoscape v3.5 (Shannon et al., 2003). The initial prior knowledge network (PKN) was generated based on known protein-protein interactions only validated experimentally. Significantly enriched pathways were analyzed within the network using CytoKEGG application within Cytoscape. The parent network and each of the individual pathway networks are available for visualization and further analysis using following cytoscape session link found within supplementary materials. Comprehensive resource of mammalian protein complexes (Corum analysis) was carried out using online tool http://mips.helmholtz-muenchen.de/corum/ (Giurgiu et al., 2018).

Functional classification GO biological process (BP), molecular function (MF) and cellular component (CC) were carried out using molecular signatures database (MSigD) within Gene Set Enrichment (GSEA) tool (Liberzon et al., 2011; Subramanian et al., 2005).

### Kaplan Meyer and Protein Expression in Clinical Samples

Analysis of the overall survival of breast cancer patients using a Kaplan-Meier plot were carried out using KMPlotter (Gyorffy et al., 2010), https://kmplot.com/analysis/ n=3951. All patients’ samples were included in the analysis shown, i.e ER, PR status, subtype, lymph node status and grade. Analysis of protein expression levels in breast tumour versus normal samples were representative of those within the Human Protein Atlas database (Uhlen et al., 2015) http://www.proteinatlas.org.

## Notes

### Competing Interest Statement

The authors have declared no competing interest.

