## Supplementary Figures for "Signalling Network of Breast Cancer Cells in Response to Progesterone"

### Fig. S1

**A**

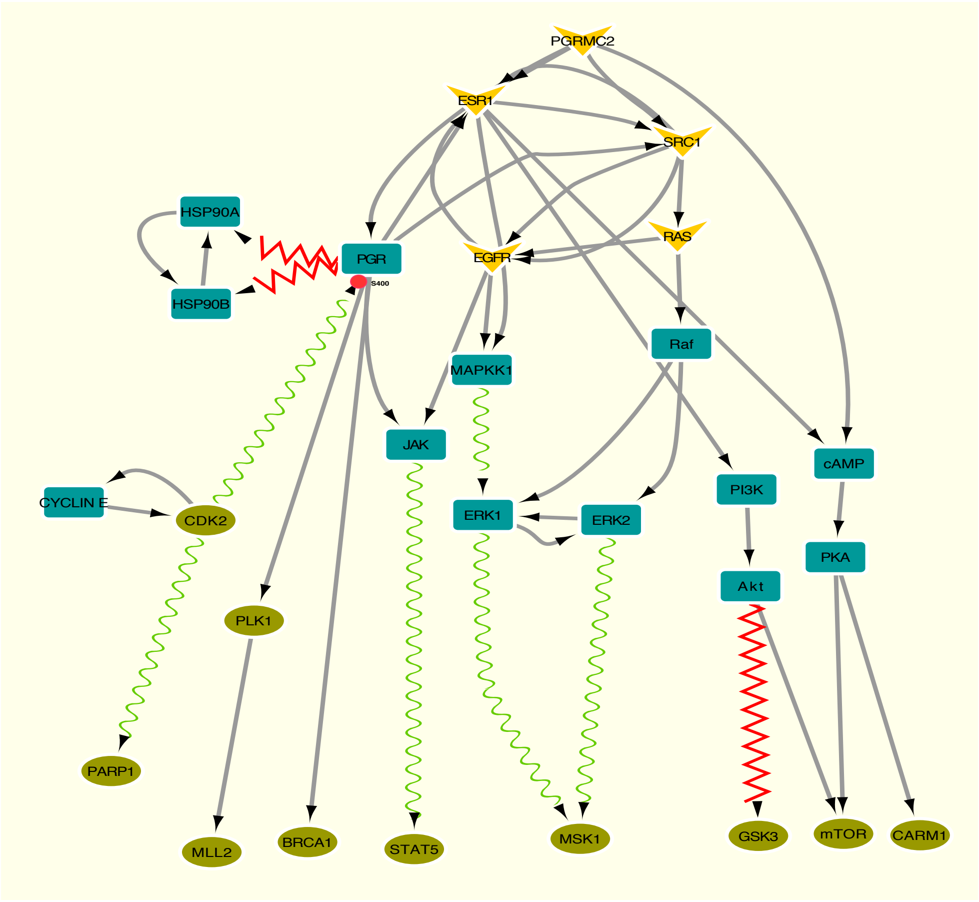


**B**

Progesterone literature

0 – 0 0 – 0

Table Panel

f(x)

| shared name | shared interaction | name | interaction | |
|---|---|---|---|---|
| CDK2 (interacts with) PGR | phosphorylates | CDK2 (interacts with) PGR | phosphorylates | Peirson 2004 Mullany |
| AKT (interacts with) GSK3 | dissociates from | AKT (interacts with) GSK3 | leaves | |
| AKT (interacts with) mTOR | interacts with | AKT (interacts with) mTOR | interacts with | |
| CDK2 (interacts with) CYCLIN E | interacts with | CDK2 (interacts with) CYCLIN E | interacts with | |
| CDK2 (interacts with) PARP1 | phosphorylates | CDK2 (interacts with) PARP1 | phosphorylates | |
| CYCLIN E (interacts with) CDK2 | interacts with | CYCLIN E (interacts with) CDK2 | interacts with | |
| EGFR (interacts with) ESR1 | interacts with | EGFR (interacts with) ESR1 | interacts with | |
| EGFR (interacts with) Jak2 | interacts with | EGFR (interacts with) Jak2 | interacts with | |
| EGFR (interacts with) MAPKK1 | interacts with | EGFR (interacts with) MAPKK1 | interacts with | |
| EGFR (interacts with) MEK | interacts with | EGFR (interacts with) MEK | interacts with | |
| ERK1 (interacts with) ERK2 | interacts with | ERK1 (interacts with) ERK2 | interacts with | |
| ERK1 (interacts with) MSK1 | phosphorylates | ERK1 (interacts with) MSK1 | phosphorylates | |
| ERK1 (interacts with) PGR | phosphorylates | ERK1 (interacts with) PGR | phosphorylates | |
| ERK2 (interacts with) ERK1 | interacts with | ERK2 (interacts with) ERK1 | interacts with | |
| ERK2 (interacts with) MSK1 | phosphorylates | ERK2 (interacts with) MSK1 | phosphorylates | |
| ESR1 (interacts with) MAPKK1 | interacts with | ESR1 (interacts with) MAPKK1 | interacts with | |
| [illegible] | [illegible] | [illegible] | [illegible] | |

**Fig. S2**

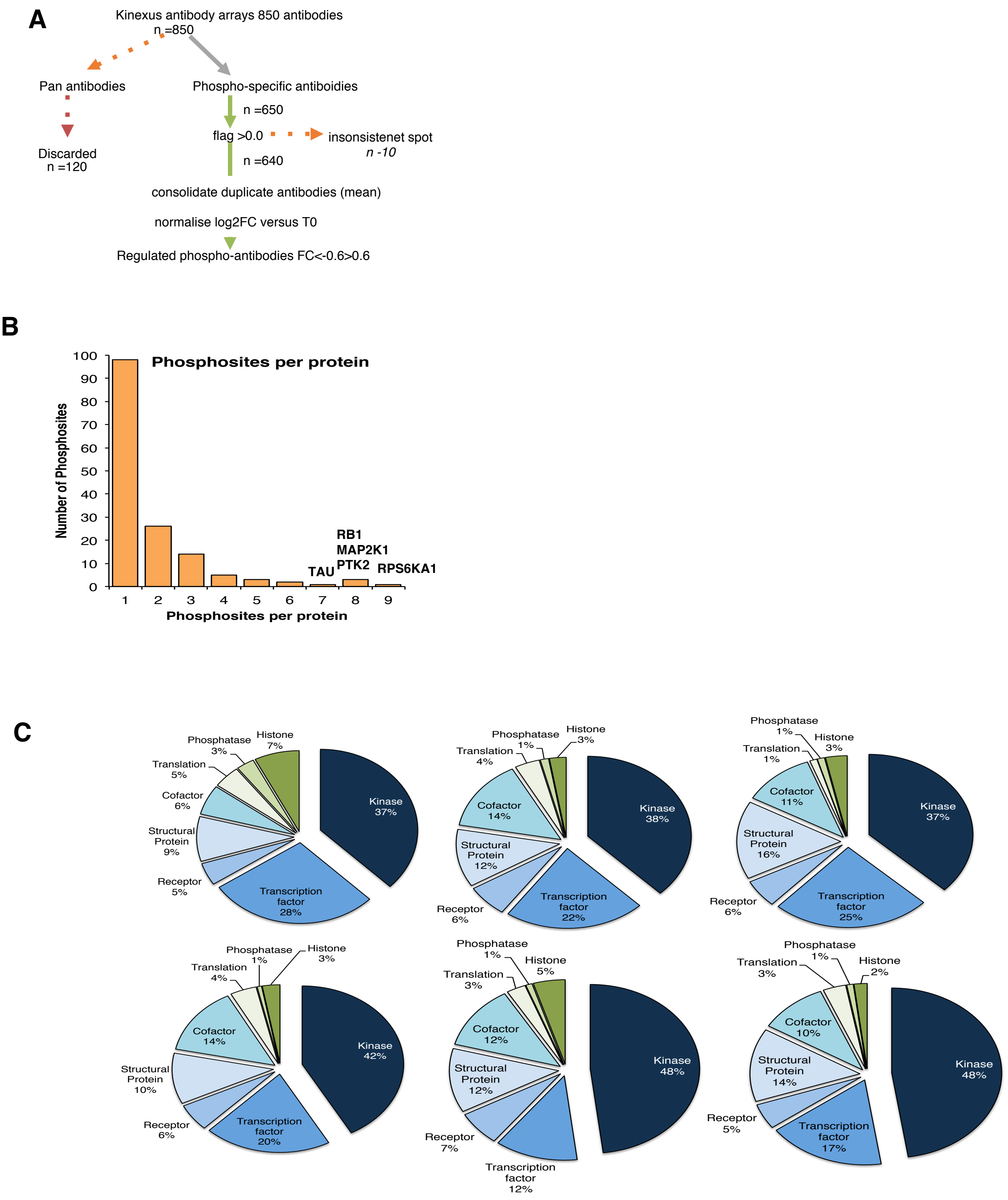
A
Kinexus antibody arrays 850 antibodies
n =850
Pan antibodies
Phospho-specific antiboidies
Discarded
n =120
n =650
flag >0.0
insonsistenet spot
n -10
n =640
consolidate duplicate antibodies (mean)
normalise log2FC versus T0
Regulated phospho-antibodies FC<-0.6>0.6
B
Phosphosites per protein
Number of Phosphosites
100
90
80
70
60
50
40
30
20
10
0
1
2
3
4
5
6
7
8
9
Phosphosites per protein
TAU
RB1
MAP2K1
PTK2
RPS6KA1
C
Histone 7%
Phosphatase 3%
Translation 5%
Cofactor 6%
Structural Protein 9%
Receptor 5%
Transcription factor 28%
Kinase 37%
Histone 3%
Phosphatase 1%
Translation 4%
Cofactor 14%
Structural Protein 12%
Receptor 6%
Transcription factor 22%
Kinase 38%
Phosphatase 1%
Histone 3%
Translation 1%
Cofactor 11%
Structural Protein 16%
Receptor 6%
Transcription factor 25%
Kinase 37%
Phosphatase 1%
Histone 3%
Translation 4%
Cofactor 14%
Structural Protein 10%
Receptor 6%
Transcription factor 20%
Kinase 42%
Phosphatase 1%
Histone 5%
Translation 3%
Cofactor 12%
Structural Protein 12%
Receptor 7%
Transcription factor 12%
Kinase 48%
Phosphatase 1%
Histone 2%
Translation 3%
Cofactor 10%
Structural Protein 14%
Receptor 5%
Transcription factor 17%
Kinase 48%

**Fig. S2 continued**

D

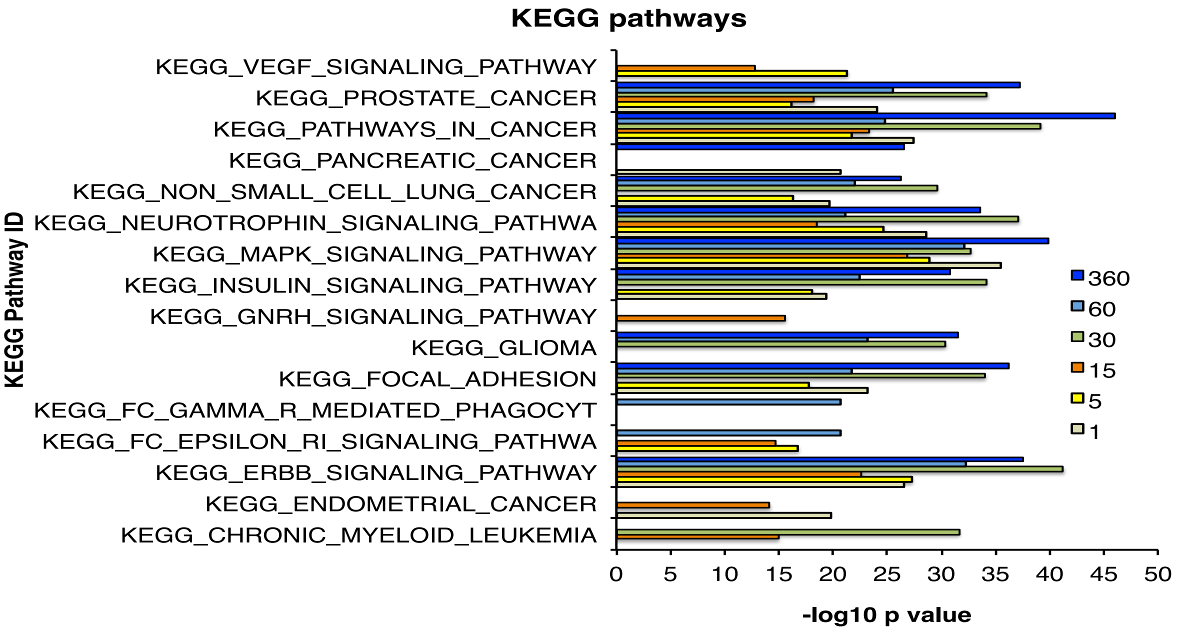


E

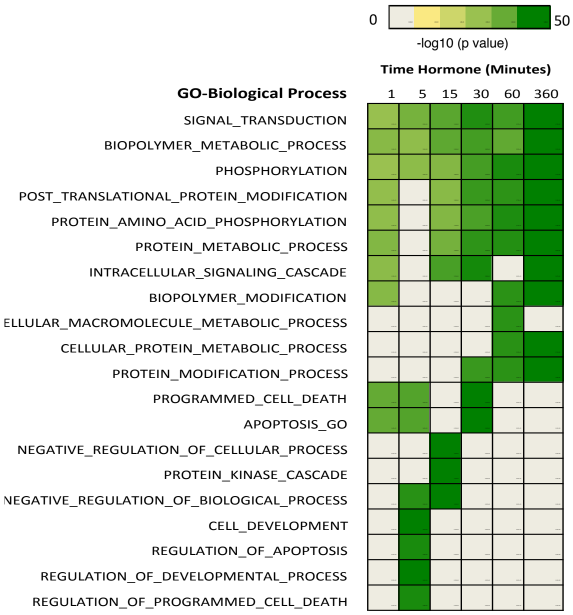


F

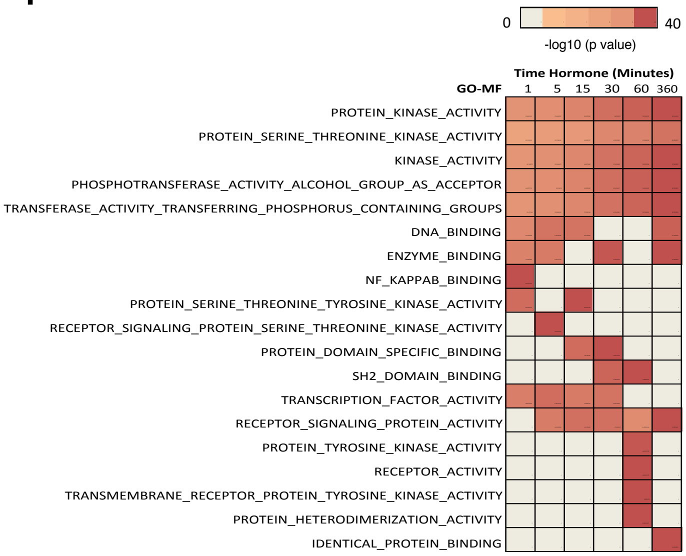

**Fig. S3**

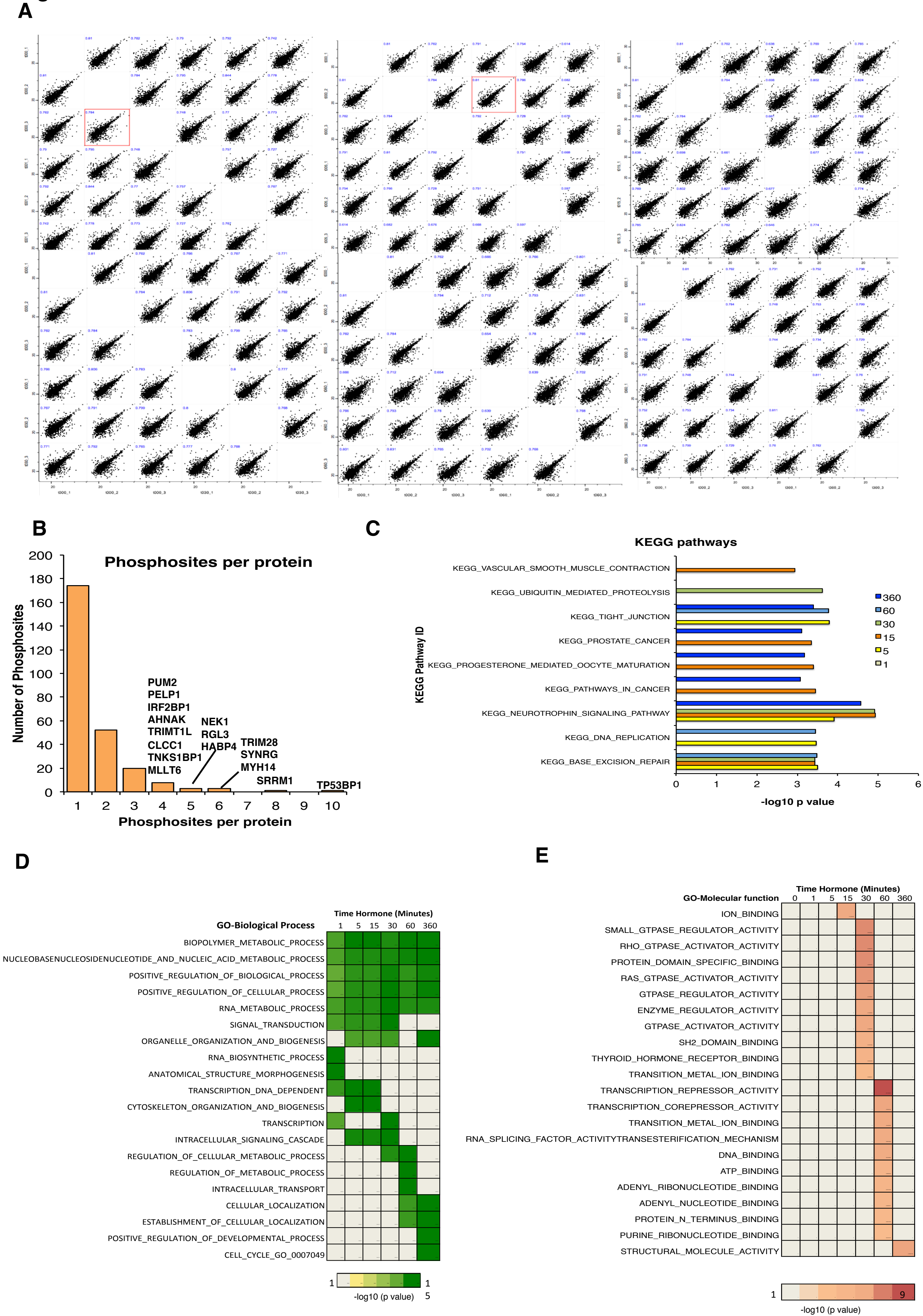
A
B
Phosphosites per protein
Number of Phosphosites
200
180
160
140
120
100
80
60
40
20
0
1 2 3 4 5 6 7 8 9 10
Phosphosites per protein
PUM2
PELP1
IRF2BP1
AHNAK
TRIMT1L
CLCC1
TNKS1BP1
MLLT6
NEK1
RGL3
HABP4
TRIM28
SYNRG
MYH14
SRRM1
TP53BP1
C
KEGG pathways
KEGG Pathway ID
KEGG_VASCULAR_SMOOTH_MUSCLE_CONTRACTION
KEGG_UBIQUITIN_MEDIATED_PROTEOLYSIS
KEGG_TIGHT_JUNCTION
KEGG_PROSTATE_CANCER
KEGG_PROGESTERONE_MEDIATED_OOCYTE_MATURATION
KEGG_PATHWAYS_IN_CANCER
KEGG_NEUROTROPHIN_SIGNALING_PATHWAY
KEGG_DNA_REPLICATION
KEGG_BASE_EXCISION_REPAIR
0 1 2 3 4 5 6
-log10 p value
360
60
30
15
5
1
D
Time Hormone (Minutes)
GO-Biological Process
1 5 15 30 60 360
BIOPOLYMER_METABOLIC_PROCESS
NUCLEOBASENUCLEOSIDENUCLEOTIDE_AND_NUCLEIC_ACID_METABOLIC_PROCESS
POSITIVE_REGULATION_OF_BIOLOGICAL_PROCESS
POSITIVE_REGULATION_OF_CELLULAR_PROCESS
RNA_METABOLIC_PROCESS
SIGNAL_TRANSDUCTION
ORGANELLE_ORGANIZATION_AND_BIOGENESIS
RNA_BIOSYNTHETIC_PROCESS
ANATOMICAL_STRUCTURE_MORPHOGENESIS
TRANSCRIPTION_DNA_DEPENDENT
CYTOSKELETON_ORGANIZATION_AND_BIOGENESIS
TRANSCRIPTION
INTRACELLULAR_SIGNALING_CASCADE
REGULATION_OF_CELLULAR_METABOLIC_PROCESS
REGULATION_OF_METABOLIC_PROCESS
INTRACELLULAR_TRANSPORT
CELLULAR_LOCALIZATION
ESTABLISHMENT_OF_CELLULAR_LOCALIZATION
POSITIVE_REGULATION_OF_DEVELOPMENTAL_PROCESS
CELL_CYCLE_GO_0007049
1
1
-log10 (p value)
5
E
Time Hormone (Minutes)
GO-Molecular function
0 1 5 15 30 60 360
ION_BINDING
SMALL_GTPASE_REGULATOR_ACTIVITY
RHO_GTPASE_ACTIVATOR_ACTIVITY
PROTEIN_DOMAIN_SPECIFIC_BINDING
RAS_GTPASE_ACTIVATOR_ACTIVITY
GTPASE_REGULATOR_ACTIVITY
ENZYME_REGULATOR_ACTIVITY
GTPASE_ACTIVATOR_ACTIVITY
SH2_DOMAIN_BINDING
THYROID_HORMONE_RECEPTOR_BINDING
TRANSITION_METAL_ION_BINDING
TRANSCRIPTION_REPRESSOR_ACTIVITY
TRANSCRIPTION_COREPRESSOR_ACTIVITY
TRANSITION_METAL_ION_BINDING
RNA_SPLICING_FACTOR_ACTIVITYTRANSESTERIFICATION_MECHANISM
DNA_BINDING
ATP_BINDING
ADENYL_RIBONUCLEOTIDE_BINDING
ADENYL_NUCLEOTIDE_BINDING
PROTEIN_N_TERMINUS_BINDING
PURINE_RIBONUCLEOTIDE_BINDING
STRUCTURAL_MOLECULE_ACTIVITY
1
9
-log10 (p value)

**Fig. S4.**

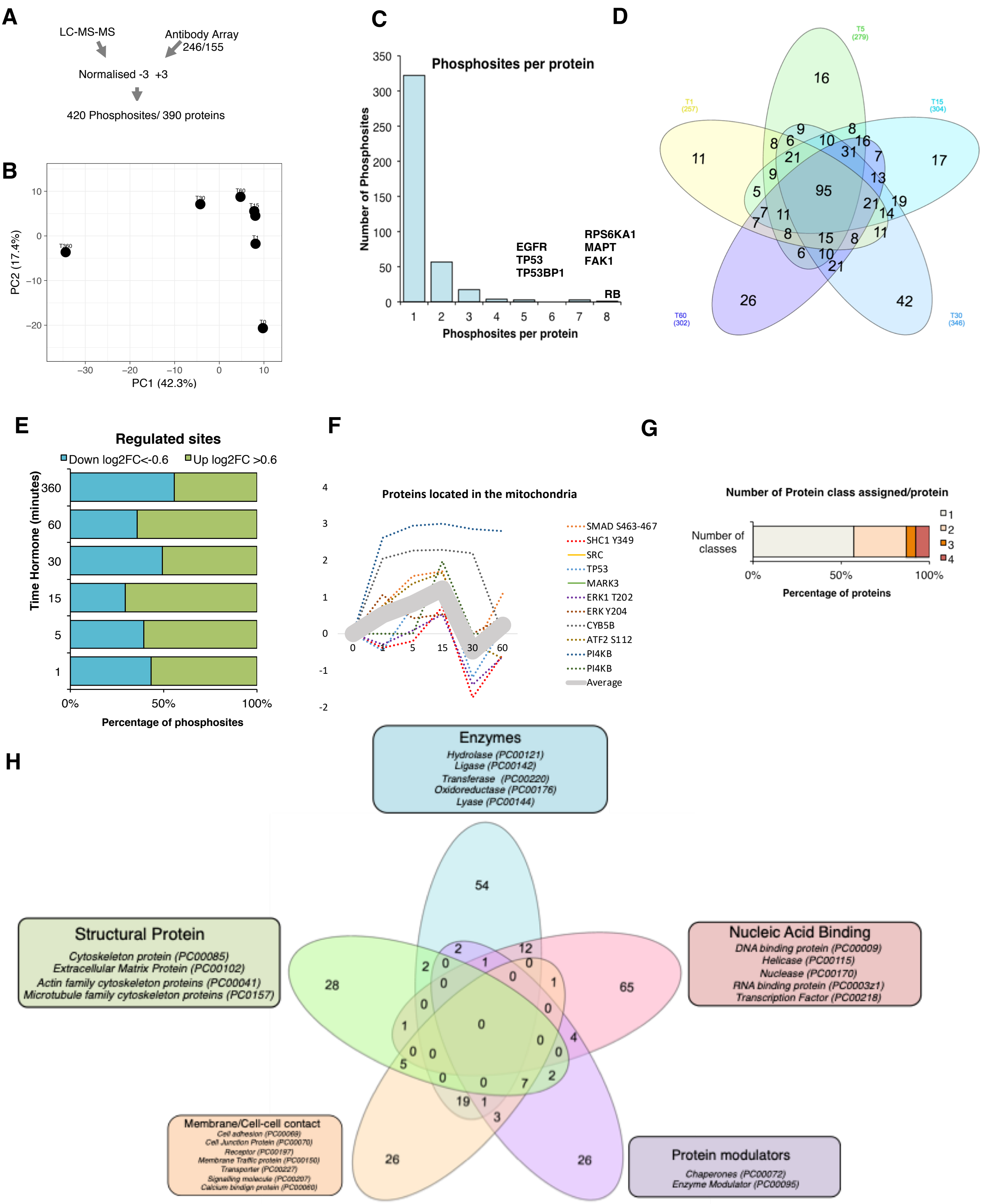
A
LC-MS-MS
Antibody Array
246/155
Normalised -3 +3
420 Phosphosites/ 390 proteins
B
T360
T30
T60
T15
T1
T0
PC2 (17.4%)
PC1 (42.3%)
C
Phosphosites per protein
Number of Phosphosites
EGFR
TP53
TP53BP1
RPS6KA1
MAPT
FAK1
RB
Phosphosites per protein
D
T5 (279)
T1 (257)
T15 (304)
T60 (302)
T30 (346)
E
Regulated sites
Down log2FC<-0.6
Up log2FC >0.6
Time Hormone (minutes)
Percentage of phosphosites
F
Proteins located in the mitochondria
SMAD S463-467
SHC1 Y349
SRC
TP53
MARK3
ERK1 T202
ERK Y204
CYB5B
ATF2 S112
PI4KB
PI4KB
Average
G
Number of Protein class assigned/protein
Number of classes
Percentage of proteins
H
Enzymes
Hydrolase (PC00121)
Ligase (PC00142)
Transferase (PC00220)
Oxidoreductase (PC00176)
Lyase (PC00144)
Structural Protein
Cytoskeleton protein (PC00085)
Extracellular Matrix Protein (PC00102)
Actin family cytoskeleton proteins (PC00041)
Microtubule family cytoskeleton proteins (PC0157)
Nucleic Acid Binding
DNA binding protein (PC00009)
Helicase (PC00115)
Nuclease (PC00170)
RNA binding protein (PC0003z1)
Transcription Factor (PC00218)
Membrane/Cell-cell contact
Cell adhesion (PC00069)
Cell Junction Protein (PC00070)
Receptor (PC00197)
Membrane Traffic protein (PC00150)
Transporter (PC00227)
Signaling molecule (PC00207)
Calcium bindign protein (PC00060)
Protein modulators
Chaperones (PC00072)
Enzyme Modulator (PC00095)

Fig. S5

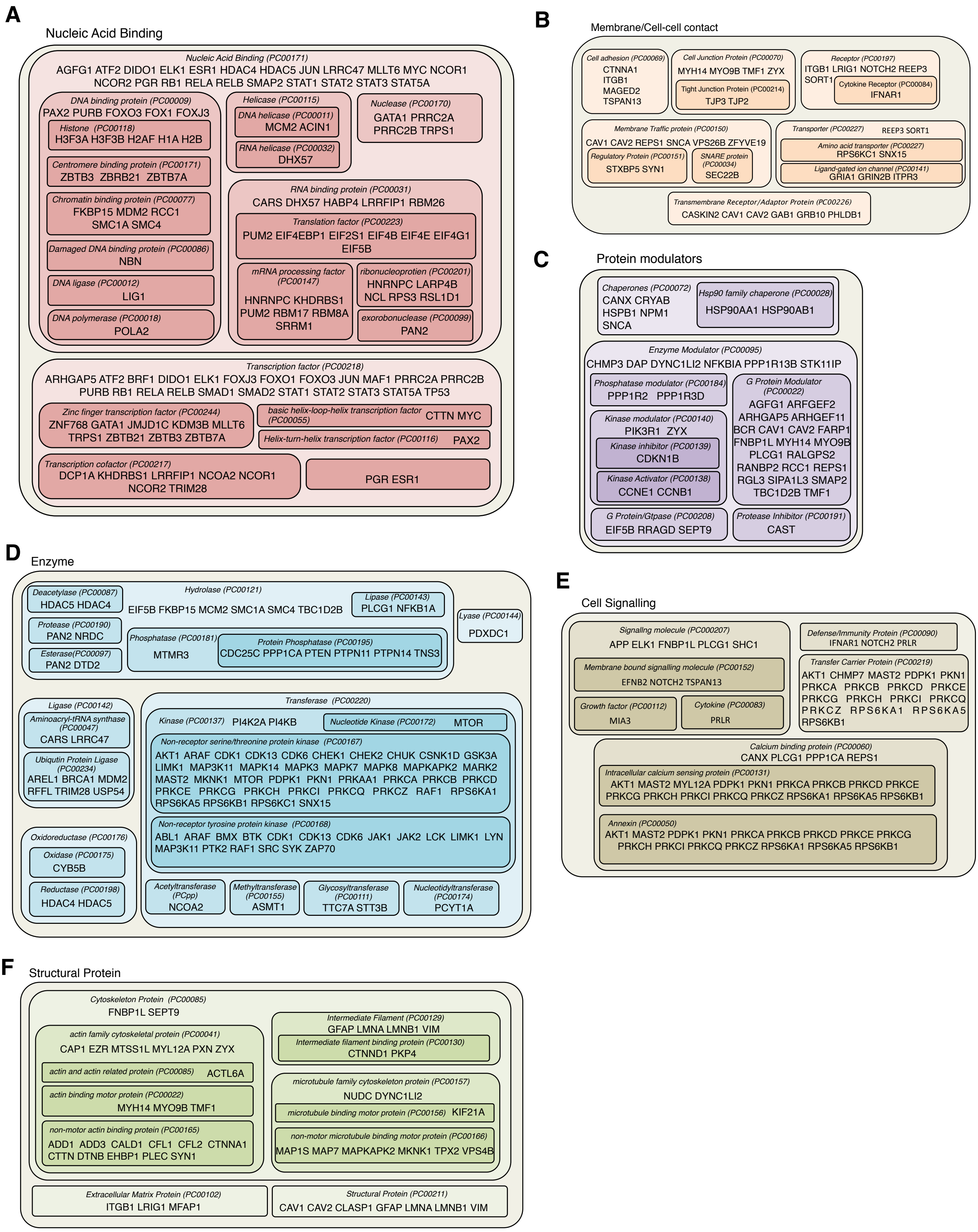
A
Nucleic Acid Binding
Nucleic Acid Binding (PC00171)
AGFG1 ATF2 DIDO1 ELK1 ESR1 HDAC4 HDAC5 JUN LRRC47 MLLT6 MYC NCOR1 NCOR2 PGR RB1 RELA RELB SMAP2 STAT1 STAT2 STAT3 STAT5A
DNA binding protein (PC00009)
PAX2 PURB FOXO3 FOX1 FOXJ3
Histone (PC00118)
H3F3A H3F3B H2AF H1A H2B
Centromere binding protein (PC00171)
ZBTB3 ZBRB21 ZBTB7A
Chromatin binding protein (PC00077)
FKBP15 MDM2 RCC1 SMC1A SMC4
Damaged DNA binding protein (PC00086)
NBN
DNA ligase (PC00012)
LIG1
DNA polymerase (PC00018)
POLA2
Helicase (PC00115)
DNA helicase (PC00011)
MCM2 ACIN1
RNA helicase (PC00032)
DHX57
Nuclease (PC00170)
GATA1 PRRC2A PRRC2B TRPS1
RNA binding protein (PC00031)
CARS DHX57 HABP4 LRRFIP1 RBM26
Translation factor (PC00223)
PUM2 EIF4EBP1 EIF2S1 EIF4B EIF4E EIF4G1 EIF5B
mRNA processing factor (PC00147)
HNRNPC KHDRBS1 PUM2 RBM17 RBM8A SRRM1
ribonucleoprotien (PC00201)
HNRNPC LARP4B NCL RPS3 RSL1D1
exorobonuclease (PC00099)
PAN2
Transcription factor (PC00218)
ARHGAP5 ATF2 BRF1 DIDO1 ELK1 FOXJ3 FOXO1 FOXO3 JUN MAF1 PRRC2A PRRC2B PURB RB1 RELA RELB SMAD1 SMAD2 STAT1 STAT2 STAT3 STAT5A TP53
Zinc finger transcription factor (PC00244)
ZNF768 GATA1 JMJD1C KDM3B MLLT6 TRPS1 ZBTB21 ZBTB3 ZBTB7A
basic helix-loop-helix transcription factor (PC00055)
CTTN MYC
Helix-turn-helix transcription factor (PC00116)
PAX2
Transcription cofactor (PC00217)
DCP1A KHDRBS1 LRRFIP1 NCOA2 NCOR1 NCOR2 TRIM28
PGR ESR1
B
Membrane/Cell-cell contact
Cell adhesion (PC00069)
CTNNA1 ITGB1 MAGED2 TSPAN13
Cell Junction Protein (PC00070)
MYH14 MYO9B TMF1 ZYX
Tight Junction Protein (PC00214)
TJP3 TJP2
Receptor (PC00197)
ITGB1 LRIG1 NOTCH2 REEP3 SORT1
Cytokine Receptor (PC00084)
IFNAR1
Membrane Traffic protein (PC00150)
CAV1 CAV2 REPS1 SNCA VPS26B ZFYVE19
Regulatory Protein (PC00151)
STXBP5 SYN1
SNARE protein (PC00034)
SEC22B
Transporter (PC00227)
REEP3 SORT1
Amino acid transporter (PC00227)
RPS6KC1 SNX15
Ligand-gated ion channel (PC00141)
GRIA1 GRIN2B ITPR3
Transmembrane Receptor/Adaptor Protein (PC00226)
CASKIN2 CAV1 CAV2 GAB1 GRB10 PHLDB1
C
Protein modulators
Chaperones (PC00072)
CANX CRYAB HSPB1 NPM1 SNCA
Hsp90 family chaperone (PC00028)
HSP90AA1 HSP90AB1
Enzyme Modulator (PC00095)
CHMP3 DAP DYNC1LI2 NFKBIA PPP1R13B STK11IP
Phosphatase modulator (PC00184)
PPP1R2 PPP1R3D
Kinase modulator (PC00140)
PIK3R1 ZYX
Kinase inhibitor (PC00139)
CDKN1B
Kinase Activator (PC00138)
CCNE1 CCNB1
G Protein Modulator (PC00022)
AGFG1 ARFGEF2 ARHGAP5 ARHGEF11 BCR CAV1 CAV2 FARP1 FNBP1L MYH14 MYO9B PLCG1 RALGPS2 RANBP2 RCC1 REPS1 RGL3 SIPA1L3 SMAP2 TBC1D2B TMF1
G Protein/Gtpase (PC00208)
EIF5B RRAGD SEPT9
Protease Inhibitor (PC00191)
CAST
D
Enzyme
Deacetylase (PC00087)
HDAC5 HDAC4
Hydrolase (PC00121)
EIF5B FKBP15 MCM2 SMC1A SMC4 TBC1D2B
Lipase (PC00143)
PLCG1 NFKB1A
Protease (PC00190)
PAN2 NRDC
Esterase(PC00097)
PAN2 DTD2
Phosphatase (PC00181)
MTMR3
Protein Phosphatase (PC00195)
CDC25C PPP1CA PTEN PTPN11 PTPN14 TNS3
Lyase (PC00144)
PDXDC1
Ligase (PC00142)
Aminoacryl-tRNA synthase (PC00047)
CARS LRRC47
Ubiqutin Protein Ligase (PC00234)
AREL1 BRCA1 MDM2 RFFL TRIM28 USP54
Oxidoreductase (PC00176)
Oxidase (PC00175)
CYB5B
Reductase (PC00198)
HDAC4 HDAC5
Transferase (PC00220)
Kinase (PC00137) PI4K2A PI4KB
Nucleotide Kinase (PC00172) MTOR
Non-receptor serine/threonine protein kinase (PC00167)
AKT1 ARAF CDK1 CDK13 CDK6 CHEK1 CHEK2 CHUK CSNK1D GSK3A LIMK1 MAP3K11 MAPK14 MAPK3 MAPK7 MAPK8 MAPKAPK2 MARK2 MAST2 MKNK1 MTOR PDPK1 PKN1 PRKAA1 PRKCA PRKCB PRKCD PRKCE PRKCG PRKCH PRKCI PRKCQ PRKCZ RAF1 RPS6KA1 RPS6KA5 RPS6KB1 RPS6KC1 SNX15
Non-receptor tyrosine protein kinase (PC00168)
ABL1 ARAF BMX BTK CDK1 CDK13 CDK6 JAK1 JAK2 LCK LIMK1 LYN MAP3K11 PTK2 RAF1 SRC SYK ZAP70
Acetyltransferase (PCpp)
NCOA2
Methyltransferase (PC00155)
ASMT1
Glycosyltransferase (PC00111)
TTC7A STT3B
Nucleotidyltransferase (PC00174)
PCYT1A
E
Cell Signalling
Signallng molecule (PC000207)
APP ELK1 FNBP1L PLCG1 SHC1
Membrane bound signalling molecule (PC00152)
EFNB2 NOTCH2 TSPAN13
Growth factor (PC00112)
MIA3
Cytokine (PC00083)
PRLR
Defense/Immunity Protein (PC00090)
IFNAR1 NOTCH2 PRLR
Transfer Carrier Protein (PC00219)
AKT1 CHMP7 MAST2 PDPK1 PKN1 PRKCA PRKCB PRKCD PRKCE PRKCG PRKCH PRKCI PRKCQ PRKCZ RPS6KA1 RPS6KA5 RPS6KB1
Calcium binding protein (PC00060)
CANX PLCG1 PPP1CA REPS1
Intracellular calcium sensing protein (PC00131)
AKT1 MAST2 MYL12A PDPK1 PKN1 PRKCA PRKCB PRKCD PRKCE PRKCG PRKCH PRKCI PRKCQ PRKCZ RPS6KA1 RPS6KA5 RPS6KB1
Annexin (PC00050)
AKT1 MAST2 PDPK1 PKN1 PRKCA PRKCB PRKCD PRKCE PRKCG PRKCH PRKCI PRKCQ PRKCZ RPS6KA1 RPS6KA5 RPS6KB1
F
Structural Protein
Cytoskeleton Protein (PC00085)
FNBP1L SEPT9
actin family cytoskeletal protein (PC00041)
CAP1 EZR MTSS1L MYL12A PXN ZYX
actin and actin related protein (PC00085) ACTL6A
actin binding motor protein (PC00022)
MYH14 MYO9B TMF1
non-motor actin binding protein (PC00165)
ADD1 ADD3 CALD1 CFL1 CFL2 CTNNA1 CTTN DTNB EHBP1 PLEC SYN1
Intermediate Filament (PC00129)
GFAP LMNA LMNB1 VIM
Intermediate filament binding protein (PC00130)
CTNND1 PKP4
microtubule family cytoskeleton protein (PC00157)
NUDC DYNC1LI2
microtubule binding motor protein (PC00156) KIF21A
non-motor microtubule binding motor protein (PC00166)
MAP1S MAP7 MAPKAPK2 MKNK1 TPX2 VPS4B
Extracellular Matrix Protein (PC00102)
ITGB1 LRIG1 MFAP1
Structural Protein (PC00211)
CAV1 CAV2 CLASP1 GFAP LMNA LMNB1 VIM

**Fig. S6**

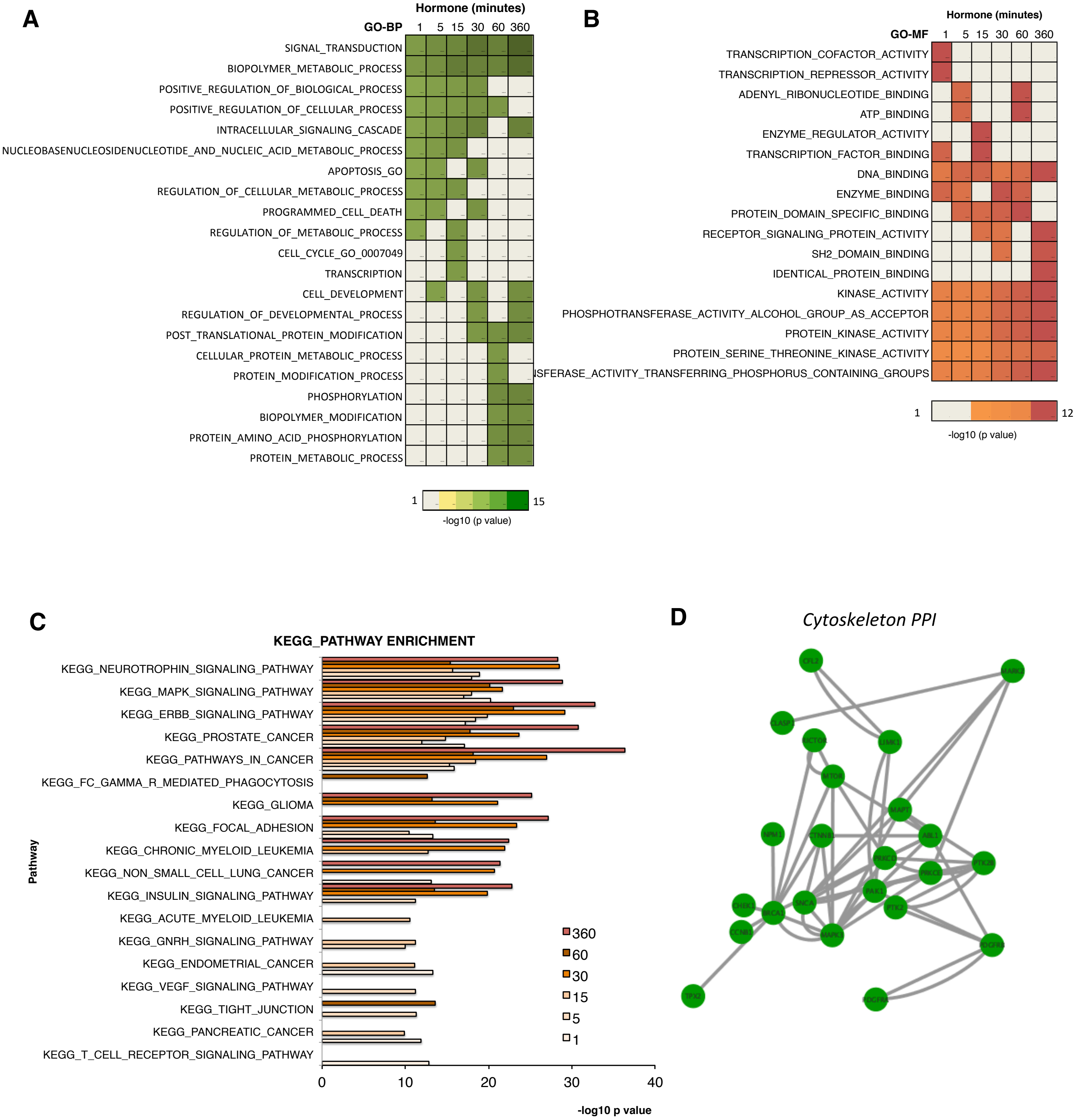
A
Hormone (minutes)
GO-BP
1
5
15
30
60
360
SIGNAL_TRANSDUCTION
BIOPOLYMER_METABOLIC_PROCESS
POSITIVE_REGULATION_OF_BIOLOGICAL_PROCESS
POSITIVE_REGULATION_OF_CELLULAR_PROCESS
INTRACELLULAR_SIGNALING_CASCADE
NUCLEOBASENUCLEOSIDENUCLEOTIDE_AND_NUCLEIC_ACID_METABOLIC_PROCESS
APOPTOSIS_GO
REGULATION_OF_CELLULAR_METABOLIC_PROCESS
PROGRAMMED_CELL_DEATH
REGULATION_OF_METABOLIC_PROCESS
CELL_CYCLE_GO_0007049
TRANSCRIPTION
CELL_DEVELOPMENT
REGULATION_OF_DEVELOPMENTAL_PROCESS
POST_TRANSLATIONAL_PROTEIN_MODIFICATION
CELLULAR_PROTEIN_METABOLIC_PROCESS
PROTEIN_MODIFICATION_PROCESS
PHOSPHORYLATION
BIOPOLYMER_MODIFICATION
PROTEIN_AMINO_ACID_PHOSPHORYLATION
PROTEIN_METABOLIC_PROCESS
1
15
-log10 (p value)
B
Hormone (minutes)
GO-MF
1
5
15
30
60
360
TRANSCRIPTION_COFACTOR_ACTIVITY
TRANSCRIPTION_REPRESSOR_ACTIVITY
ADENYL_RIBONUCLEOTIDE_BINDING
ATP_BINDING
ENZYME_REGULATOR_ACTIVITY
TRANSCRIPTION_FACTOR_BINDING
DNA_BINDING
ENZYME_BINDING
PROTEIN_DOMAIN_SPECIFIC_BINDING
RECEPTOR_SIGNALING_PROTEIN_ACTIVITY
SH2_DOMAIN_BINDING
IDENTICAL_PROTEIN_BINDING
KINASE_ACTIVITY
PHOSPHOTRANSFERASE_ACTIVITY_ALCOHOL_GROUP_AS_ACCEPTOR
PROTEIN_KINASE_ACTIVITY
PROTEIN_SERINE_THREONINE_KINASE_ACTIVITY
ISFERASE_ACTIVITY_TRANSFERRING_PHOSPHORUS_CONTAINING_GROUPS
1
12
-log10 (p value)
C
KEGG_PATHWAY ENRICHMENT
Pathway
KEGG_NEUROTROPHIN_SIGNALING_PATHWAY
KEGG_MAPK_SIGNALING_PATHWAY
KEGG_ERBB_SIGNALING_PATHWAY
KEGG_PROSTATE_CANCER
KEGG_PATHWAYS_IN_CANCER
KEGG_FC_GAMMA_R_MEDIATED_PHAGOCYTOSIS
KEGG_GLIOMA
KEGG_FOCAL_ADHESION
KEGG_CHRONIC_MYELOID_LEUKEMIA
KEGG_NON_SMALL_CELL_LUNG_CANCER
KEGG_INSULIN_SIGNALING_PATHWAY
KEGG_ACUTE_MYELOID_LEUKEMIA
KEGG_GNRH_SIGNALING_PATHWAY
KEGG_ENDOMETRIAL_CANCER
KEGG_VEGF_SIGNALING_PATHWAY
KEGG_TIGHT_JUNCTION
KEGG_PANCREATIC_CANCER
KEGG_T_CELL_RECEPTOR_SIGNALING_PATHWAY
360
60
30
15
5
1
0
10
20
30
40
-log10 p value
D
Cytoskeleton PPI

**Fig. S7**

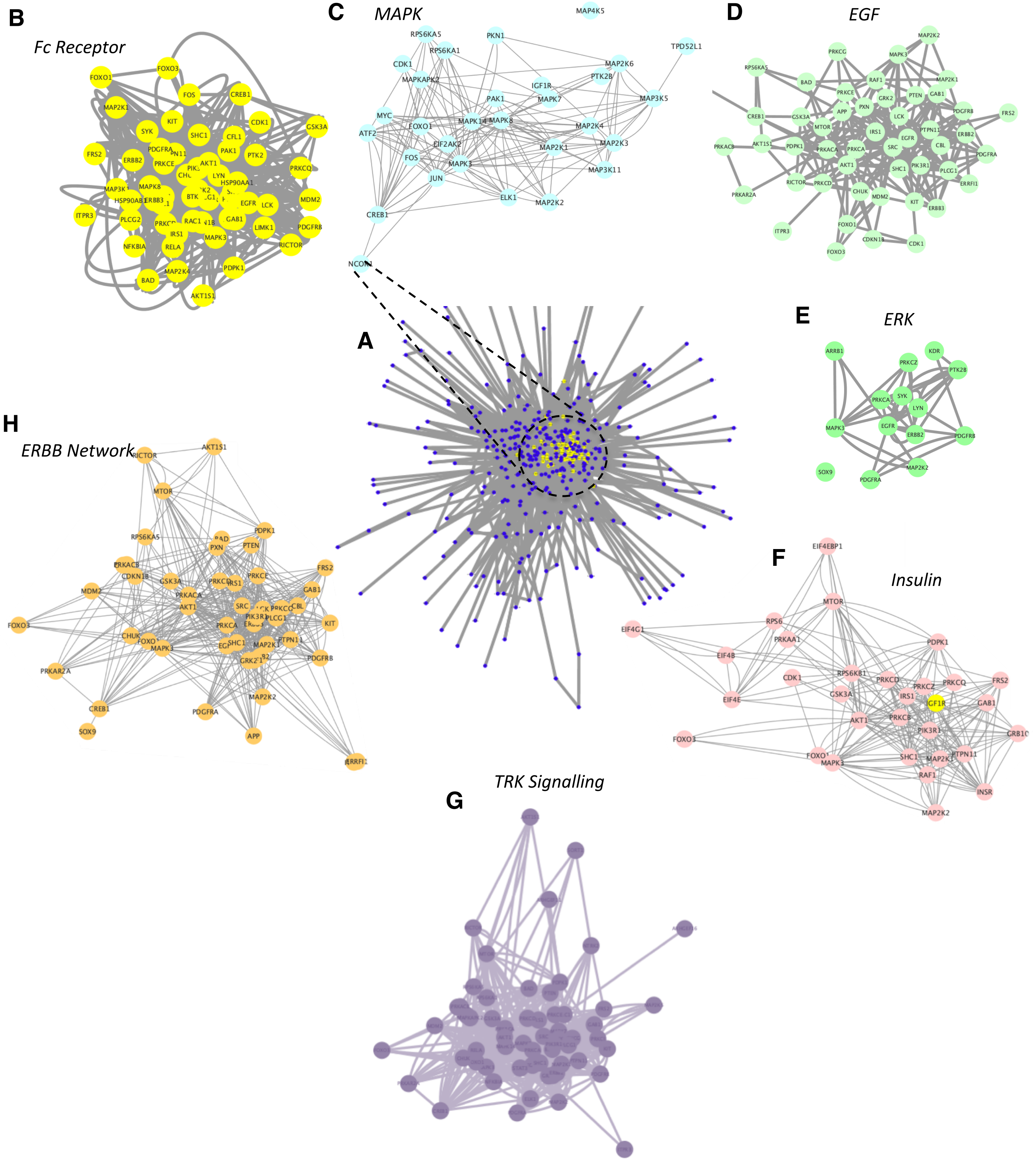
B
Fc Receptor
FOXO1
FOXO3
FOS
CREB1
MAP2K1
KIT
CDK1
GSK3A
SYK
SHC1
CFL1
FRS2
ERBB2
PAK1
PTK2
PRKCE
PRKCQ
MAP3K1
MAPK8
HSP90AA1
HSP90AB1
ERBB3
BTK
EGFR
LCK
MDM2
ITPR3
PLCG2
RAC1
GAB1
LIMK1
PDGFRB
IRS1
NFKBIA
RELA
MAPK3
RICTOR
MAP2K4
PDPK1
BAD
AKT1S1
AKT1
LYN
C
MAPK
MAP4K5
RPS6KA5
PKN1
RPS6KA1
TPD52L1
CDK1
MAP2K6
MAPKAPK2
PTK2B
IGF1R
PAK1
MAPK7
MAP3K5
MYC
ATF2
FOXO1
MAPK14
MAPK8
MAP2K4
EIF2AK2
MAP2K3
FOS
MAPK3
MAP2K1
MAP3K11
JUN
ELK1
MAP2K2
CREB1
NCOR1
D
EGF
MAP2K2
PRKCG
MAPK3
RPS6KA5
BAD
RAF1
MAP2K1
PRKCE
PXN
GRK2
PTEN
GAB1
CREB1
GSK3A
APP
LCK
PDGFRB
FRS2
MTOR
IRS1
EGFR
PTPN11
ERBB2
PRKACB
AKT1S1
PDPK1
PRKACA
PRKCA
SRC
CBL
PDGFRA
AKT1
SHC1
PIK3R1
PLCG1
RICTOR
PRKCD
CHUK
MDM2
ERRFI1
PRKAR2A
KIT
ERBB3
FOXO1
ITPR3
CDKN1B
CDK1
FOXO3
A
E
ERK
ARRB1
KDR
PRKCZ
PTK2B
SYK
PRKCA
LYN
MAPK3
EGFR
ERBB2
PDGFRB
SOX9
MAP2K2
PDGFRA
H
ERBB Network
RICTOR
AKT1S1
MTOR
PDPK1
RPS6KA5
BAD
PXN
PTEN
PRKACB
CDKN1B
GSK3A
PRKCD
IRS1
PRKCE
FRS2
MDM2
PRKACA
GAB1
AKT1
SRC
PRKCQ
CBL
PIK3R1
PLCG1
FOXO3
PRKCA
KIT
CHUK
FOXO1
SHC1
MAP2K1
PTPN11
MAPK3
PDGFRB
PRKAR2A
GRK2
MAP2K2
CREB1
PDGFRA
SOX9
APP
ERRFI1
F
Insulin
EIF4EBP1
MTOR
RPS6
EIF4G1
PRKAA1
PDPK1
EIF4B
CDK1
RPS6KB1
PRKCD
PRKCZ
PRKCQ
FRS2
GSK3A
IRS1
IGF1R
GAB1
EIF4E
AKT1
PRKCB
PIK3R1
GRB10
FOXO3
FOXO1
MAPK3
SHC1
MAP2K1
PTPN11
RAF1
INSR
MAP2K2
G
TRK Signalling
